# Within-species floral odor variation is maintained by spatial and temporal heterogeneity in pollinator communities

**DOI:** 10.1101/2020.09.29.318089

**Authors:** Mark A. Szenteczki, Adrienne L. Godschalx, Andrea Galmán, Anahí Espíndola, Marc Gibernau, Nadir Alvarez, Sergio Rasmann

## Abstract

Floral odor is a complex trait that mediates many biotic interactions, including pollination. While high intraspecific floral odor variation appears to be common, the ecological and evolutionary drivers of this variation are often unclear. Here, we investigated the influence of spatially and temporally heterogeneous pollinator communities on floral odor variation in *Arum maculatum* (Araceae). Through Europe-wide field surveys, we identified high floral odor diversity and shifts in the dominant pollinator species within several *A. maculatum* populations compared to pollinator data from the same sites ten years ago. Using common-garden experiments, we further confirmed that inflorescences from native and foreign pollinator backgrounds were equally efficient at attracting local pollinators. The substantial within-population floral odor variation we observed may therefore be advantageous when facing temporally heterogeneous pollinator communities. We propose spatio-temporal heterogeneity in pollinators as one potential mechanism maintaining diverse floral odor bouquets in angiosperms.

## INTRODUCTION

The rapid diversification of angiosperms relative to other land plants has long fascinated evolutionary biologists, dating back to Darwin (1903). Today, we can attribute much of the variation in angiosperm diversification rates to plant-pollinator interactions (Hernández-Hernández and Wiens 2020). As the dominant pollinators across all terrestrial ecosystems, insects play a key role in floral trait evolution (Sprengel 1793, Henslow 1888; Grant 1949; Raven 1977; Kevan and Baker 1983; Crepet and Niklas 2009; van der Niet and Johnson 2012; Schiestl and Johnson 2013). These plant-insect interactions are mediated by multimodal floral signals (reviewed in Junker and Parachnowitsch 2015), which include odor (Holopainen 2004; Raguso 2008ab; Whitehead and Peakall 2009), color (Goyret et al. 2007; du Plessis et al. 2018), morphology (Ayasse et al. 2003; Ibanez et al. 2010), nectar (Parachnowitsch et al. 2019), and pollen (Dobson and Bergström 2000). Nowadays, we are able to identify many plant volatile organic compounds (VOCs) with relative ease (Knudsen and Gershenzon 2020). However, the ecological and evolutionary drivers of floral odor variation often remain unclear, due to difficulties in collecting and integrating data on floral odor variation and spatio-temporal variation in pollinators.

In a recent review, Delle-Vedove et al. (2017) found that intraspecific floral odor variation is frequently identified within diverse angiosperm lineages. Since heterogenous pollinator communities are known to influence variation in many plant traits (Schemske and Horvitz 1988; Fishbein and Venable 1996; Price et al. 2005; Weber et al. 2019), it can be argued that floral odor variation should also be influenced by spatio-temporal variation in pollinators (Delle-Vedove et al. 2017, Haveramp et al. 2018, Dormont et al. 2019, Friberg et al. 2019, Burkle et al. 2020, Farré-Armengol et al. 2020). However, only a small fraction of studies to date have been able to explicitly link floral odor variation and variation in pollinator community structure (e.g. Chess et al. 2008; Klahre et al. 2011; Breitkopf et al. 2013; Peter and Johnson 2014; Sun et al. 2013; Gross et al. 2016). Several factors may explain this discrepancy, including phylogenetic constraints in the biosynthetic pathways for the production of VOCs (Raguso et al. 2006), tradeoffs between pollinator attraction and chemical defense against herbivory (Schiestl et al. 2014), biochemical and energetic limitations (Delle-Vedove et al. 2011), genetic drift (Suinyuy et al. 2012), or gene flow (Svensson et al. 2005). Moreover, most of the aforementioned surveys of floral odor and pollinator community variation were carried out at a single timepoint. As a result, the impact of temporal heterogeneity in pollinator communities on floral odor variation remains unknown; here, we aim to address this gap in our knowledge.

If pollinator communities vary temporally within populations, heterogeneous and/or frequency-dependent selective pressures may favor the maintenance of floral odor variation through balancing selection (Delph and Kelly 2013; Wittmann et al. 2017; Bertram and Masel 2019). Alternatively, if pollinators only vary spatially among populations, we might expect divergent selective pressures to occur, leading to local adaptation (Leimu and Fischer 2008, Gómez et al. 2009; Gross et al. 2016; Gervasi and Schiestl 2017; Suinyuy and Johnson 2018; Sayers et al. 2020). Variation may also result from phenotypic plasticity under variable abiotic conditions or biotic interactions (Majetic et al. 2009; Campbell et al. 2019), if several floral odor phenotypes can result from a single genotype. Here, we investigated how each of these three processes influence floral odor variation, using wild populations of *Arum maculatum* (Araceae).

Several aspects of *A. maculatum*’s ecology make it a suitable model system for studying the relationship between pollinators and floral odor. *Arum maculatum* inflorescences attract and temporarily trap their pollinators, mainly the moth flies *Psychoda phalaenoides* and *Psycha grisescens* (Diptera: Psychodidae), using complex blends of VOCs which appear to vary across the species distribution range (Kite 1995; Kite et al. 1998; Diaz and Kite 2002; Chartier et al. 2011; Chartier et al. 2013). These odor blends are thought to mimic the odor of dung or rotting organic matter, the natural brood sites of these Psychodidae (Lack and Diaz 1991). Pollinators are trapped in a highly specialized floral chamber (Bröderbauer et al. 2013) until the day after VOC emissions (Gibernau et al. 2004). As a result, we are able to accurately quantify the complete pollinator community attracted by each inflorescence, and unambiguously relate pollinator abundances to VOC emissions. Since *A. maculatum* inflorescences do not offer any rewards to their pollinators (Lack and Diaz 1991), and Psychodidae appear to be attracted by VOCs alone (Dormer 1960; Kite 1995; Urru et al. 2011), the reproductive success of an individual inflorescence (i.e. its ability to attract efficient pollinators) should be closely tied to its unique floral odor. Selfing is also avoided through protogyny; female flowers are no longer receptive to pollen when the male flowers mature (Lack and Diaz 1991).

This unique pollination ecology provides an opportunity to address key questions on how and why floral odor variation is maintained in widely distributed species. Namely, is variation maintained across species ranges, due to spatial variation in pollinators, or within populations, due to temporally heterogeneous pollinators? Spatial and temporal variation in pollinators may drive floral odor variation through one or more of the aforementioned evolutionary processes (i.e. balancing selection, local adaptation, or plasticity). Below, we summarize the patterns in floral odor diversity we expected to observe under each scenario:

1) Balancing selection is expected to maintain greater variation in deceptively pollinated species compared to rewarding species (Ackerman et al. 2011), through temporal variation in pollinator communities (Schemske and Horvitz 1989), frequency-dependent selection driven by pollinator learning (Ayasse et al. 2000), or relaxed selection (Salzmann et al. 2007). The tetraploid genome of *A. maculatum* (Turco et al. 2014) may allow balancing selection to maintain variation within individual genomes as well (Mable et al. 2018). If *A. maculatum* floral odor is a balanced polymorphic trait, we would expect to observe high variation in floral odor both within and among populations. This variation may be a consequence of either: a) temporally heterogeneous pollinator communities within populations, whose varying preferences maintain floral odor diversity, or b) pollinator learning, if pollinator communities are stable within populations through time.

2) Populations of *A. maculatum* are known to predominantly attract either *P. phalaenoides* or *P. grisescens* in different regions across their distribution (Espíndola et al. 2010). Following the ‘local vs. foreign’ definition of local adaptation (Kawecki and Ebert 2004) we might alternatively predict that *A. maculatum* populations harbor distinct floral odor variations, which maximize the attraction of their dominant local pollinator species. If these variations are true local adaptations, we would expect to observe decreased pollinator attraction efficiency when *A. maculatum* inflorescences are transplanted to non-native pollinator backgrounds.

3) As organisms rooted in place, plants benefit from adaptive plasticity in response to variable environmental conditions or biotic interactions (Majetic et al. 2009; Metlen et al. 2009). If *A. maculatum* floral odor is a plastic trait, we would expect to observe a correlation between climatic variables and floral odor variation across the wide geographic range of *A. maculatum*, and/or a shift in VOC emissions when inflorescences are transplanted to non-native environments.

Here, we tested these three sets of predictions by surveying natural variation in *A. maculatum* floral odor and pollinator attraction across Europe, and by transplanting individuals from all sampled populations to two common garden sites dominated by different pollinator species (either *Psychoda phalaenoides* or *Psycha grisescens*). Importantly, we selected sampling sites with pollinator data collected approximately ten years ago by Espíndola et al. (2010), allowing us to investigate temporal heterogeneity in pollinator communities within our study. Together, these data allowed us to understand the extent of within- and between-population variation in *A. maculatum* floral scent, how this variation is organized across a wide geographic range, and how both spatial and temporal pollinator community dynamics influence floral odor variation.

## MATERIALS AND METHODS

### Field sampling sites

We sampled eleven populations of *A. maculatum*, including three in France (Forêt du Gâvre, Conteville, and Chaumont), two in Switzerland (Neuchâtel and Cortaillod), two in Italy (Montese and Rifreddo), one in Croatia (Visuć), two in Serbia (Gostilje and Sokobanja), and one in Bulgaria (Chiflik), as shown in Appendix S1, Figure S1. Full sampling information, including location data and sample sizes are given in Appendix S1, Table S1. We selected Forêt du Gâvre and Neuchâtel, which are respectively dominated by *Psycha grisescens* and *Psychoda phalaenoides*, as our two common garden sites. Five to ten *A. maculatum* individuals from each population were potted and transplanted to both sites; inflorescences from Forêt du Gâvre and Neuchâtel were also reciprocally transplanted as part of this experiment.

### Floral odor and pollinator sampling

During the typical flowering period of *A. maculatum* (April – May), we conducted: 1) field sampling between 2017 and 2019, and 2) common garden experiments in Neuchâtel in 2018 and 2019, and in Forêt du Gâvre in 2019. In both field surveys and common garden experiments, we collected dynamic headspace VOCs and identified pollinators using identical methods, as detailed in Appendix S2: Supplementary Methods. Briefly, we collected VOCs from *A. maculatum* inflorescences undergoing anthesis in the early evening on polydimethylsiloxane (PDMS) coated Twister® stir bars (Gerstel: Mülheim an der Ruhr, Germany), at a rate of 200 mL min^−1^ for 30 minutes. Twisters® were kept on ice in sealed glass containers until gas chromatography–mass spectrometry (GC-MS) analysis, where volatiles were thermally desorbed and separated on a HP-5MS column. During the morning following VOC sampling, we collected all insects trapped within inflorescences and preserved them in 70% ethanol until identification.

### Testing for balancing selection and temporal heterogeneity in pollinators

First, we investigated Europe-wide patterns in floral odor variation, by calculating Bray-Curtis similarities between all individuals in our *in situ* field surveys, and visualizing the result using nonmetric multidimensional scaling (NMDS). We then assessed the effect of population (fixed effect factor) on the entire VOCs matrix using permutational multivariate analysis of variance (PERMANOVA, Bray-Curtis distance, n = 999 permutations) with the *adonis* function in the R v.3.6.1 (R Core Team 2019) package *vegan* (Oksanen et al. 2019). Then, we compared the extent of both within- and between-population variation in floral odor using Bray-Curtis similarity matrices, visualizing the resulting similarity scores using boxplots.

Second, we investigated shifts in the dominant pollinators trapped by *A. maculatum* over the past decade, both in terms of 1) Psychodidae species and 2) other insect families, comparing our observations with data (mean abundances) from the same sites in Espíndola et al. (2010). We began by calculating the mean quantities of each pollinator species trapped per inflorescence within each population. Then, to investigate shifts in pollinators, we calculated Bray-Curtis distances between pollinator communities now (2017-2019) and approximately a decade ago (2006-2008) for all sites with data from both timepoints, and visualized these results using NMDS ordinations.

### Testing for local adaptation to pollinators using common garden experiments

First, we visualized geographic patterns in mean pollinator attraction for each population *in situ*, and following transplants to both common garden sites. Then, following the ‘local vs. foreign’ definition of local adaptation (Kawecki and Ebert 2004), we tested the hypotheses that: in the Neuchâtel common garden, transplanted *A. maculatum* inflorescences which attract *P. phalaenoides* in their native population should catch 1) more *P. phalaenoides* and/or 2) more insects in total than inflorescences which attract *P. grisescens* in their native population. We expected the opposite pattern in the Forêt du Gâvre common garden (i.e. transplanted inflorescences that attract *P. grisescens* in their native population should perform better on average in Forêt du Gâvre). These predictions were tested using a two-way ANOVA on log+1 transformed pollinator counts, including ‘native pollinator’ (i.e. *P. phalaenoides*- or *P. grisescens*-dominated) and ‘common garden location’ (i.e. whether the common garden site was dominated by *P. phalaenoides* or *P. grisescens*) and their interaction as fixed factors. Here, a significant interaction would suggest local adaptation to native pollinator communities. Finally, we plotted ‘deme × habitat’ interactions (per Kawecki and Ebert 2004)—deme and habitat referring respectively to a local population and its local environmental conditions (here the pollinator community)—for mean attraction rates of *P. phalaenoides, P. grisescens*, and all insects. Data for these plots were subset based on each inflorescence’s native pollinator and common garden location, as described above.

### Testing for phenotypic plasticity and climatic correlates of floral odor variation

To address the correlation between climatic variation across Europe, floral odor, and pollinator communities trapped within inflorescences, we used two complementary approaches. First, we clustered individuals based on similarities in their proportional VOC emissions using unsupervised learning algorithms (as detailed in Appendix S2: Supplementary Methods) to test for VOC bouquet convergence due to spatial segregation or local adaptation to pollinators (Andersson et al. 2002; Hetherington-Rauth and Ramírez 2016). We then investigated whether the resulting floral odor clusters were differentially attractive to pollinator groups, including Psychodidae (identified to the species level), Brachycera, Nematocera, and Staphylinidae. To control for potential biases caused by differences in the dominant pollinator species among all sampled populations, we split these comparisons based on whether individuals were sampled in a site typically dominated by *P. phalaenoides* or *P. grisescens*. We used Kruskal-Wallis tests to determine whether clusters attracted different pollinator species in each pollinator background.

Next, we investigated the influence of large-scale climatic variation on floral odor variation, by performing a mantel test (Euclidean distance, 999 replicates) between 19 bioclimatic layers (BIO1-BIO19) at 30-seconds resolution from WorldClim (Hijmans et al. 2005) and population-level VOC data from our field surveys. Then, we investigated whether the types or proportions of odor variations within populations shifted as result of transplanting by comparing the relative abundances of each VOC cluster, both within populations of origin and following transplants.

Finally, we investigated whether specific VOCs were associated with the attraction of *P. phalaenoides* or *P. grisescens*. We identified candidate compounds using the Random Forest implementation in the R package *randomForest* (Liaw and Wiener 2002), with permutation importance enabled (*ntree* = 500, *mtry* = 8; optimized using the *tuneRF* function). Then, we calculated conditional feature contributions, and identified the combinations of compounds which had the greatest influence on the predictive strength of the model above, using the Python package *TreeInterpreter* (Saabas 2019). Finally, we performed Mann-Whitney *U*-tests to identify shifts in the candidate compounds most strongly associated with *P. phalaenoides* or *P. grisescens* attraction. Specifically, we tested whether populations with sufficiently large sample sizes (*n* ≥8) emitted different quantities of candidate compounds, when comparing between the two common gardens.

## RESULTS

### Balancing selection and temporal heterogeneity in pollinators

We observed that *A. maculatum* floral odor is highly variable across Europe. After filtering out compounds present in blank samples, we retained 18 *A. maculatum* floral VOCs present in relative abundances above 1% (Table 1). Many of the major compounds we identified (e.g. indole, p-cresol, 2-heptanone, β-citronellene, and three unnamed sesquiterpenes) have been previously reported in studies of *A. maculatum* floral odor (Diaz and Kite 2002; Chartier et al. 2013; Marotz-Clausen et al. 2018). Populations across Europe differed in their proportional emissions of VOCs *in situ* (PERMANOVA, R^2^ = 0.38, Pr(>F) = 0.001). However, we observed substantial within- and among-population variation in floral odor (Appendix S1, Figure S2), resulting in no clear regional differentiation in the NMDS ordination (Appendix S1, Figure S3).

**Table 1.** Proportional VOC blend compositions of 147 *Arum maculatum* individuals from 11 populations. The average blends for both K-medoids (PAM) clusters identified in this study are shown below, and visualized in Appendix S1, Figure S7.

| Compound | K1 (%)<br>N=97 | K2 (%)<br>N=50 |
| --- | --- | --- |
| 2-heptanone | 1.84 | 1.19 |
| β-citronellene | 4.31 | 2.09 |
| cis β-ocimene | 0.47 | 0.54 |
| p-cresol | 1.68 | 0.94 |
| **indole \*\*\*** | **24.35** | **65.39** |
| **skatole \*\*\*** | **2.52** | **1.03** |
| α-copaene | 0.14 | 0.76 |
| **Z-caryophyllene \*** | **8.37** | **3.28** |
| **β-caryophyllene \*** | **2.59** | **0.83** |
| α-humulene | 6.68 | 2.92 |
| alloaromadendrene | 8.13 | 3.45 |
| β-humulene | 3.54 | 1.87 |
| α-selinene | 0.44 | 0.33 |
| bicyclogermacrene | 7.76 | 2.44 |
| d-cadinene | 2.11 | 0.83 |
| Unnamed sesquiterpene<br>(RI 1404) | 4.97 | 2.06 |
| **Unnamed sesquiterpene**<br>**(RI 1473) \*** | **5.76** | **2.37** |
| Unnamed sesquiterpene<br>(RI 1681) | 14.35 | 7.70 |

\* denotes compounds that vary significantly between groups (Kruskal-Wallis test, $p < 0.05$ , $df = 1$ )

We also observed that *A. maculatum* pollinator communities are temporally variable. Ultimately, we were able to re-survey pollinators trapped by *A. maculatum* inflorescences in six populations (listed in Figure 1), approximately 10 years after the surveys conducted by Espíndola et al. (2010). The total insect communities trapped by *A. maculatum* appear to have shifted in most of these populations -only Forêt du Gâvre remained relatively consistent over the past decade (Figure 1). When focusing on Psychodidae only, we observed shifts in the dominant psychodid pollinator trapped within inflorescences in four out of six populations: Chaumont FR, Conteville FR, Gostilje SRB, and Visuć, HR (Figure 1). Additionally, we identified a substantial decrease in the average quantity of pollinators trapped within inflorescences in Conteville FR: from 47.6 *P. phalaenoides* and 2.3 *P. grisescens* per inflorescence in Espíndola et al. (2010), to 0.4 *P. phalaenoides* and 1.3 *P. grisescens* in our sampling (full results in Appendix S1, Table S2).

**Figure 1.**
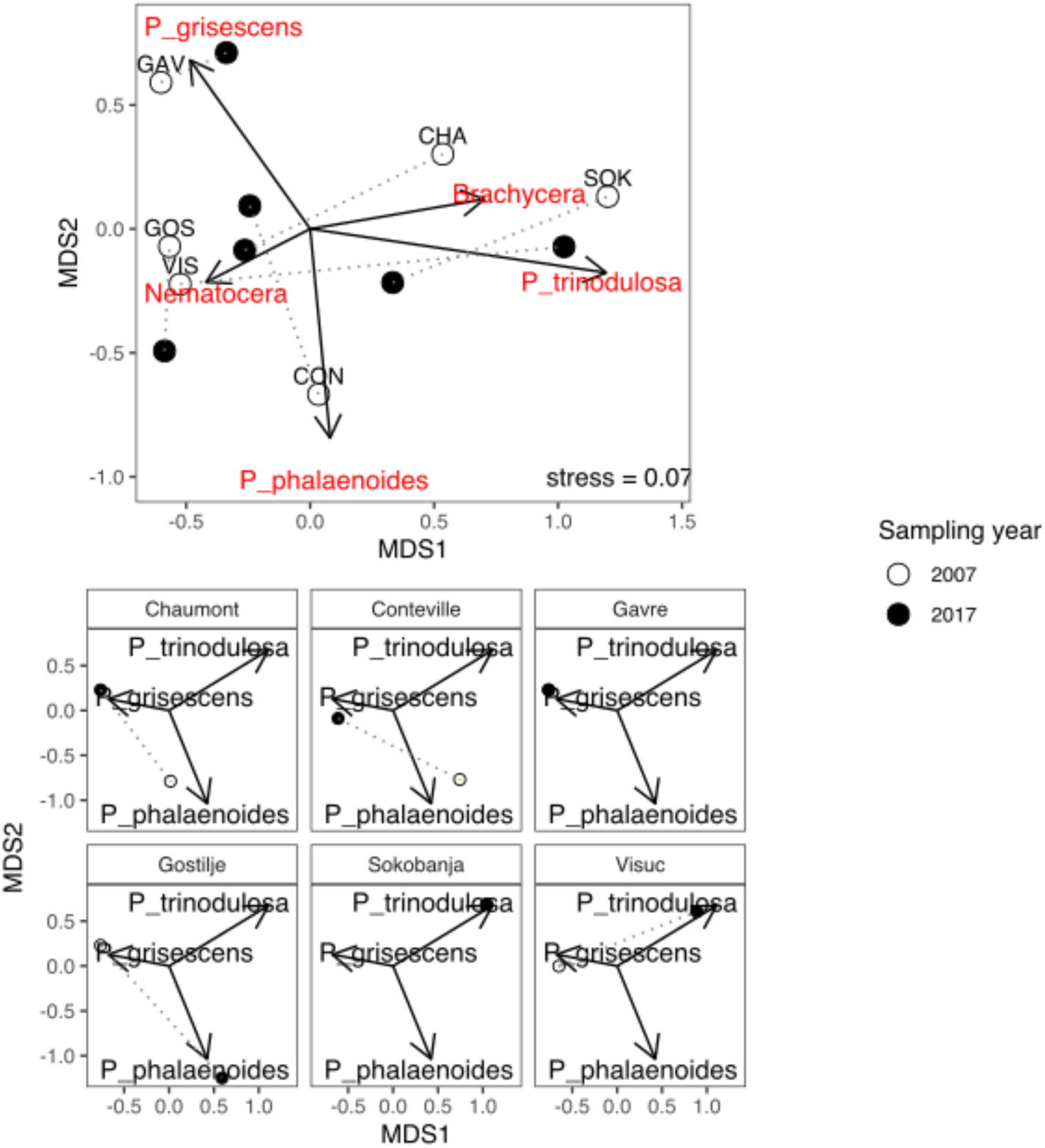
Shifts in the total pollinator communities (upper plot) and Psychodidae species (lower plots) trapped by *Arum maculatum* inflorescences. Over the past decade, the dominant Psychodid pollinator appears to have shifted in four out of six re-sampled populations.

### Local adaptation to pollinators

We observed that transplanted inflorescences typically attracted the dominant local pollinator species as efficiently as native inflorescences in both common gardens (Figure 2). While common garden location had a significant effect on the quantity of *P. phalaenoides* and *P. grisescens* caught by *A. maculatum* (i.e. the two transplant sites were indeed dominated by different Psychodidae species), no native pollinator × common garden location interaction effect was observed (2-way ANOVA, Pr(>F) > 0.05; full results in Appendix S1, Table S3). Together, these results show that *A. maculatum* populations are generally not locally adapted to a single pollinator species (Figure 3), with some possible exceptions in Forêt du Gâvre, and the two Serbian populations Gostilje and Sokobanja, which will be discussed below.

**Figure 2.**
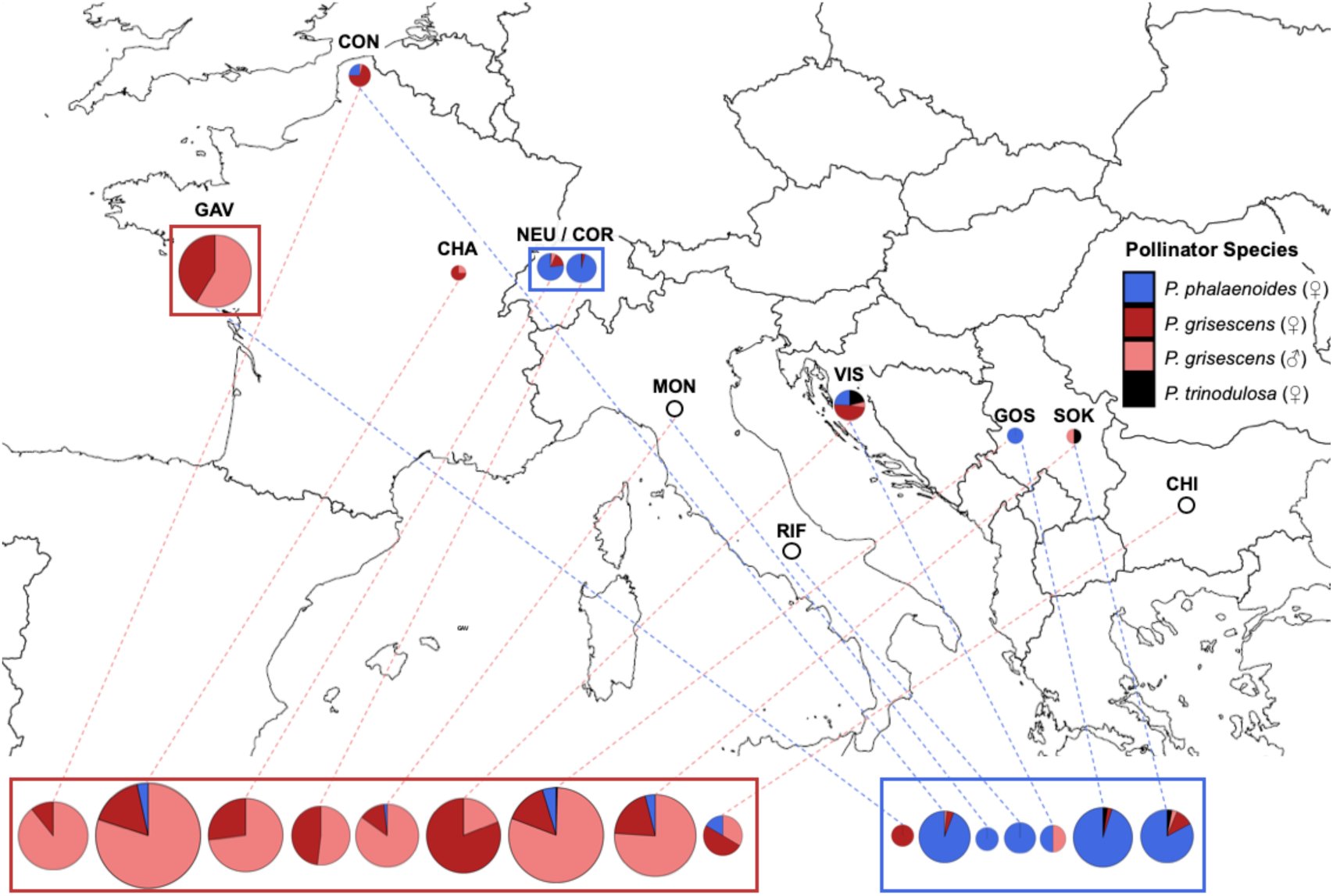
*Psychoda* spp. trapped by *Arum maculatum* inflorescences during field surveys, and following transplants to two common garden sites. Dotted lines link each field survey result (plots placed on the map) with the two corresponding transplant results (plots below map). Note: Pie charts are scaled to represent the square root +1 (to visualize small differences) of the average number of Psychodidae per plant. Empty charts represent populations where inflorescences did not attract any Psychodidae during field surveys.

**Figure 3.**
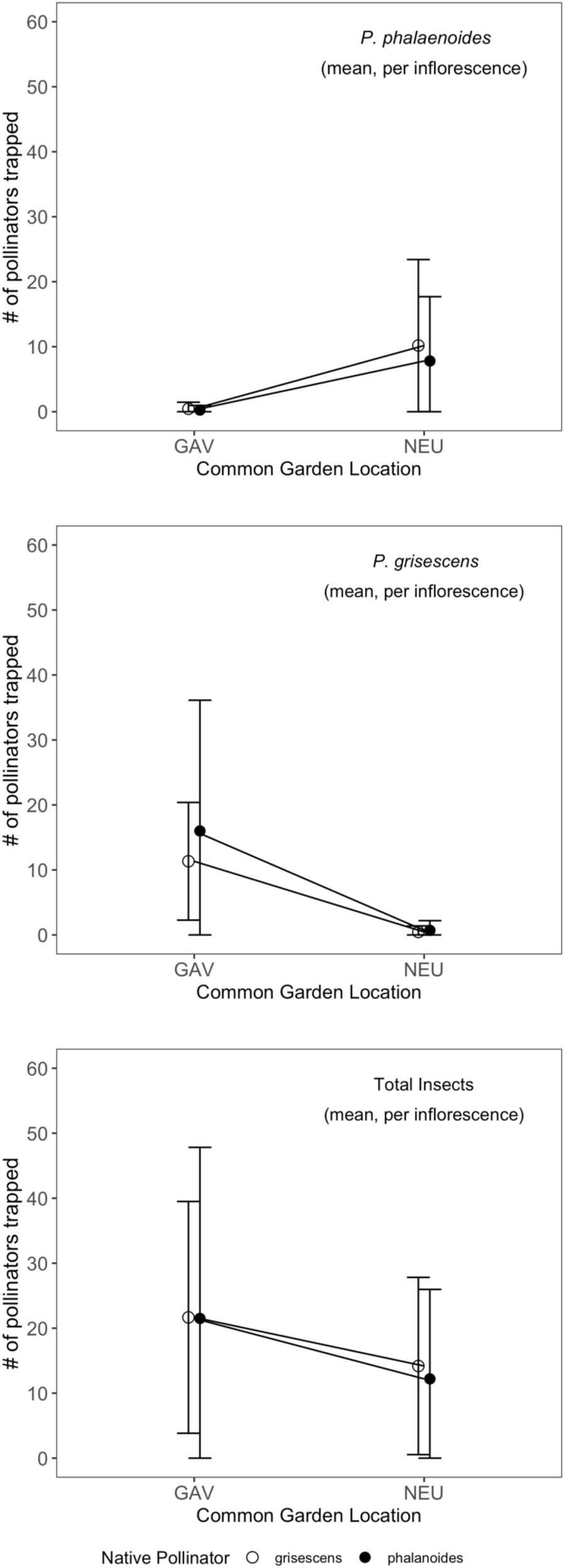
Mean (+/-SE) numbers of *Psychoda phalaenoides, Psycha grisescens*, and total insects trapped by inflorescences originating from *P. phalaenoides*-dominated sites (filled circles) and *P. grisescens-*dominated sites (hollow circles), in two common garden sites. No deme × habitat interactions (indicative of local adaptation) were identified.

While populations generally may not be locally adapted to specific pollinators, not all floral odor blends were equally attractive to all pollinator species. Notably, one population (Forêt du Gâvre) continued to exclusively attract their native pollinator *P. grisescens* when transplanted to the Neuchâtel common garden, despite *P. phalaenoides* being the most abundant species in Neuchâtel when these transplanted inflorescences opened (see Appendix S1, Figure S4). Additionally, transplanted inflorescences occasionally attracted ‘non-dominant’ Psychodidae species in both common garden sites, and a third pollinator species (*Psychoda trinodulosa*) was also identified within inflorescences in Croatia and Serbia during our field surveys (Figure 2). This species was also observed in the Neuchâtel common garden, but not in Forêt du Gâvre. Inflorescences from both Serbian populations (Gostilje and Sokobanja) continued to occasionally attract *P. trinodulosa* when transplanted the Neuchâtel common garden.

### Phenotypic plasticity and climatic correlates of floral odor variation

The cluster validation process identified two groups, clustered using the PAM (Partitioning Around Medoids) algorithm, to be optimal for our VOC dataset (Appendix S1, Figures S5 and S6). Cluster 1 was characterized by high mono- and sesquiterpene emissions, while Cluster 2 was characterized by high indole emissions. Five compounds (indole, skatole, Z-caryophyllene, β-caryophyllene, and unnamed sesquiterpene RI 1473) varied significantly between PAM clusters (Kruskal-Wallis test, *p* < 0.05, df = 2). The aforementioned unnamed sesquiterpene (Kovats index apolar 1473) has been previously identified in the floral scent of *Arum italicum* (M. Gibernau, unpublished data). The average VOC blends of both PAM clusters are summarized in Table 1, and visualized in Appendix S1, Figure S7. Both clusters were widely distributed across Europe; indole-dominated Cluster 2 was not observed in Rifreddo or Visuć, though smaller sample sizes in these populations may have contributed to this result (Appendix S1, Figure S8). Ultimately, whether an inflorescence belonged to Cluster 1 or 2 did not appear to strongly influence pollinator attraction, with considerable variation observed within both Clusters (Appendix S1, Figure S9). We did not identify any significant differences between Clusters 1 and 2, in terms of species-specific Psychodidae attraction (Kruskal-Wallis tests, *p* > 0.05, df = 2), after controlling for the dominant pollinator species where each sample was collected.

Random Forest analyses identified one compound positively correlated with *P. phalaenoides* attraction (β-humulene), and three compounds positively correlated with *P. grisescens* attraction (unnamed sesquiterpenes RI 1473 and 1681, and α-selinene). After evaluating combined feature contributions within the above Random Forest model (Appendix S1, Figure S10), we found that the predictive strength of the model was most strongly influenced by unnamed sesquiterpene (RI 1681) alone. Other strong combinations included unnamed sesquiterpene RI 1681 paired with unnamed sesquiterpene RI 1473 or α-selinene, as well as β-humulene alone, mirroring the results in our initial variable importance plot (full *Treeinterpreter* results in Appendix S1, Table S4).

We observed that inter-population variation in VOC blends overall did not correlate with bioclimatic variables (Mantel test, *p* = 0.823). Average sesquiterpene emissions remained relatively consistent between *in situ* samples and following transplants to Neuchâtel, but appeared to increase after inflorescences were transplanted to Forêt du Gâvre (Appendix S1, Figure S11). Among the four compounds linked to species-specific Psychodidae attraction, we only observed significant shifts between reciprocally transplanted inflorescences from Neuchatel and Forêt du Gâvre (Mann-Whitney tests, full result in Appendix S1, Table S5). For the two other populations with sufficiently large sample sizes (Montese, IT and Chaumont, FR), we did not observe significant differences in the emissions of these four compounds when comparing between the two common garden sites (Appendix S1, Table S5). Together, these results suggest that plasticity may contribute to the floral odor variation we observed (e.g. in the case of transplanted inflorescences from Neuchatel), but in general, the effect of transplanting on inflorescences appears to be relatively minor in comparison to natural intrapopulation variation in floral odor.

## DISCUSSION

In this study, we performed a range-wide survey of *Arum maculatum* floral odor and pollinator attraction, which identified substantial within-population variation in floral odor (Appendix S1, Figures S2 and S3), and shifts in pollinator community composition in several populations over the past decade (Figure 1). Through common garden experiments, we further demonstrated that *A. maculatum* populations typically are not locally adapted to attract exclusively *Psychoda phalaenoides* or *Psycha grisescens* (Figures 2 and 3), though local adaptation might have contributed to pollinator attraction patterns in one French and two Serbian populations. Excepting host-pathogen case studies (reviewed in Delph and Kelly 2013), our study is among the first to demonstrate that temporally fluctuating selective pressures may act to maintain high variation in a key plant trait.

Balancing selection may maintain variation within populations through several selective regimes, including relaxed selection, negative frequency-dependent selection due to pollinator learning, or environmental heterogeneity (Delph and Kelly 2013). While relaxed selection on floral odor has been observed in some angiosperms (Salzmann et al. 2007; Ibanez et al. 2010), this is unlikely the case for *A. maculatum*, given that pollinator attraction is driven by floral odor alone (Dormer 1960; Lack and Diaz 1991). Furthermore, while pollinator learning has been shown to maintain polymorphism in floral color (Gigord et al. 2001) and odor (Ayasse et al. 2000), we would argue that this is less likely to occur in the case of *A. maculatum*. Since several Psychodidae species are trapped at rates that appear to vary over short time scales, the resulting selective pressure on pollinators is likely too inconsistent to lead to adaptive pollinator learning (Renner 2006). This leaves variable conditions (i.e. pollinator communities) as the mechanism most likely contributing to the maintenance of diverse floral odor bouquets in *A. maculatum*. Further research on temporal variation in pollinators may provide greater clarity in other cases where floral trait variation cannot be explained by pollinator learning (e.g. Pellegrino et al. 2005; Jersáková et al. 2006). Our results may also explain why evidence for greater floral odor diversity in deceptive species compared to rewarding species is weak (Ackerman et al. 2011) or largely absent (Delle-Vedove et al. 2017).

Building on reciprocal transplants between two *A. maculatum* populations conducted by Chartier et al. (2013), we found that local adaptation is counteracted by temporally heterogeneous pollinator communities across Europe, with one possible exception. The Forêt du Gâvre population in France appears to have lost the ability to attract *P. phalaenoides*, either due to local adaptation resulting from the large, stable populations of *P. grisescens* we observed in their native habitat (Figure 1) or genetic drift, which is known to occur at the limits of species ranges (Geber 2011; Gould et al. 2013). Similarly, the fact that the two Serbian populations continued to attract a third psychodid species (*P. trinodulosa*) in the Neuchâtel common garden, while inflorescences from other regions did not, also argues for some level of local adaptation (Figure 2). These patterns suggest that specific compounds may also be differentially attractive to *P. trinodulosa*, but this species was too infrequently observed in our study to make any firm conclusions. These results thus do not exclude the possibility that individual VOCs are differentially attractive to certain Psychodidae species, which might be expected given previously reported differences in antennal sensilla (Faucheux and Gibernau 2011).

Through Random-Forest analyses, we identified four VOCs correlated with the attraction of *P. phalaenoides* (β-humulene) and *P. grisescens* (unidentified sesquiterpenes RI 1473 and 1681, and α-selinene). Indole, another abundant VOC in the *A. maculatum* odor bouquet, appears to be generally attractive to females of both species (Appendix S1, Figure S9 and S10), consistent with previous findings (Kite et al. 1998). Further research using Gas Chromatography -Electroantennography (Cork et al. 1990) could confirm which VOCs elicit species-specific responses, and whether these biologically active compounds are maintained at frequencies expected under balancing selection.

Adaptive plasticity is the optimal solution in situations of environmental heterogeneity when possible (Kawecki and Ebert 2004), and floral odor is known to vary with changing environmental conditions (Burkle and Runyon 2017). Recently, a “Genomic Storage Effect” has been proposed (Gulisija et al. 2016), whereby balanced polymorphism is promoted by adaptive plasticity resulting from temporally varying selection. In such a scenario, a portion of the population acts as a store of variation (i.e. compounds attractive to specific pollinators) until conditions (i.e. dominant pollinator species) change (Chesson 2000). We found that individuals from Neuchâtel and Cortaillod emitted proportionally lower quantities of indole following transplants to Forêt du Gâvre (Appendix S1, Figure S8). This result is likely due to a combination of high inter-individual variation in floral odor, and phenotypic plasticity driven by environmental variation. Furthermore, in the case of polyploids such as *A. maculatum*, the Genomic Storage Effect could even be at work at the within-individual level. Currently, it is not known whether *P. phalaenoides* and *P. grisescens* phenologies are influenced by environmental conditions; our data suggest that at least in Forêt du Gâvre, *P. phalaenoides* may emerge slightly later than *P. grisescens* (Appendix S1, Figure S4b). If environmental variation influences pollinator phenology, then plasticity in floral odor based on environmental cues could enhance pollinator attraction. Consequently, plasticity may have contributed to the variation in the four candidate compounds linked to species-specific pollinator attraction (Appendix S1, Figure S11). However, in most cases, we did not observe significant shifts consistent with plasticity (Appendix S1, Figure Table S5). After correction for multiple testing, we only found a significant shift in the emission of unnamed sesquiterpene RI 1473 (correlated with *P. grisescens* attraction) when comparing native and transplanted inflorescences from Neuchâtel. This result demonstrates that the emission of some specific compounds may be influenced environmental variation. While we cannot yet fully disentangle plasticity from variation resulting from balancing selection, the high diversity in floral odor we observed in our common garden sites is unlikely to be the result of plasticity alone.

High gene flow may also lead to the maintenance of floral odor variation. The short adult lifespan (approx. one week) and limited dispersal capacity of Psychodidae (Lack and Diaz 1991) implies that *A. maculatum* gene flow is likely driven by seed dispersal, which is mainly carried out by frugivorous birds (Snow and Snow 1988). There appears to be a strong barrier to gene flow between populations from north/central Europe and from Italy and the Balkans (Espíndola and Alvarez 2011), yet floral odor variation is shown to be widely maintained across this barrier, suggesting that — at least regionally — balancing selection driven by heterogenous pollinator community composition is at work, in association with some level of local adaptation and plasticity.

## Conclusion

While trait variation often appears to be more strongly influenced by spatial heterogeneity in selection than temporal heterogeneity (Hedrick 1986) – as demonstrated by the extensive literature on local adaptation in plants (Leimu and Fischer 2008; Anderson et al. 2011) – our study highlights how temporal heterogeneity in pollinators may also be a contributing factor in maintaining highly diverse floral odor bouquets. Although earlier models could not always demonstrate the maintenance of polymorphism through temporal heterogeneity (Hedrick 1976), recent models (e.g. Gulisija et al. 2016) provide a mechanism by which frequent shifts in pollinator communities may maintain trait variation and counteract local adaptation. To date, almost all studies have sampled floral odors and pollinators at a single timepoint — possibly contributing to the numerous cases where floral odor diversity appears to exceed pollinator diversity (Delle-Vedove et al. 2017). Further research on the temporal dynamics of pollinator communities has the potential to advance our understanding on how and why many flowering plant lineages maintain high diversity in key traits such as floral odor.

## ACKNOWLEDGEMENTS

We thank Gregory Roeder for his assistance with the processing of our VOC samples, Jérôme Albre for his assistance in Psychodidae identification, and Alberto Garcia Jimenez and Monica Fleisher for their dedicated assistance during our field sampling. We also thank Laurent Oppliger and colleagues at the Jardin Botanique de Neuchâtel for their support in hosting and maintaining our experimental populations of *Arum* inflorescences. The project was funded by the Swiss National Science Foundation through grant 31003A_163334 awarded to N.A and S.R.

## Appendix S2 SUPPLEMENTARY METHODS

**Figure S1.**
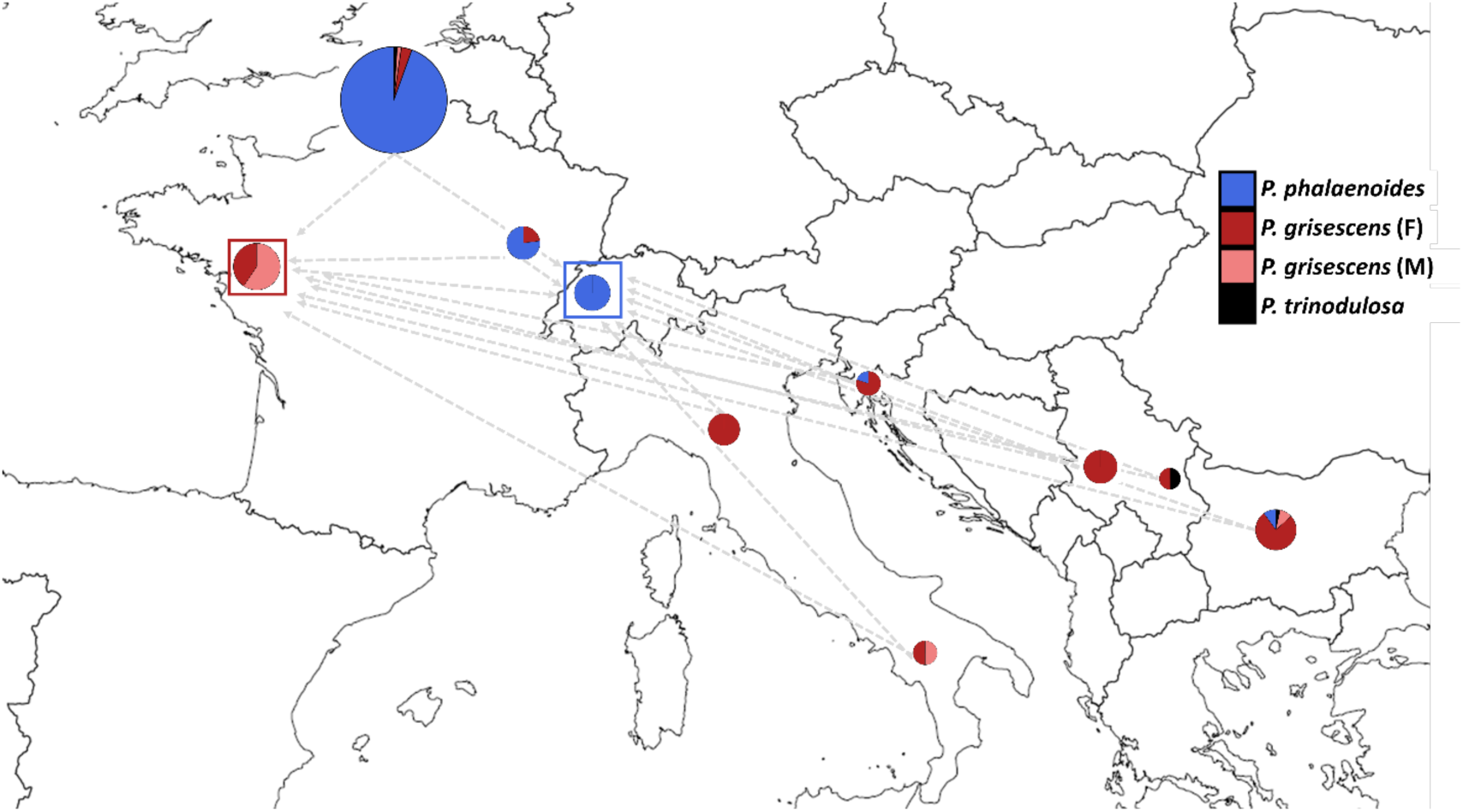
Outline of all *Arum maculatum* populations surveyed in this study, and the location of our two common garden sites (red and blue squares). Pie charts visualize data on *Psychoda* species caught by inflorescences between 2007-2009, taken from Espíndola et al. (2010). We attempted to re-survey each of these sites in this study between 2017 and 2019 (result shown in Figure 2). Note: Graph sizes represent the square root (plus one to visualize species differences in small pies) of the mean number of psychodids per plant in each population. Gray dotted arrow indicate direction for transplanting a subset of inflorescences to both common garden sites. Swiss pollinator data in this figure represents Lausanne, replaced by Neuchâtel and Cortaillod populations in the present study.

**Figure S2.**
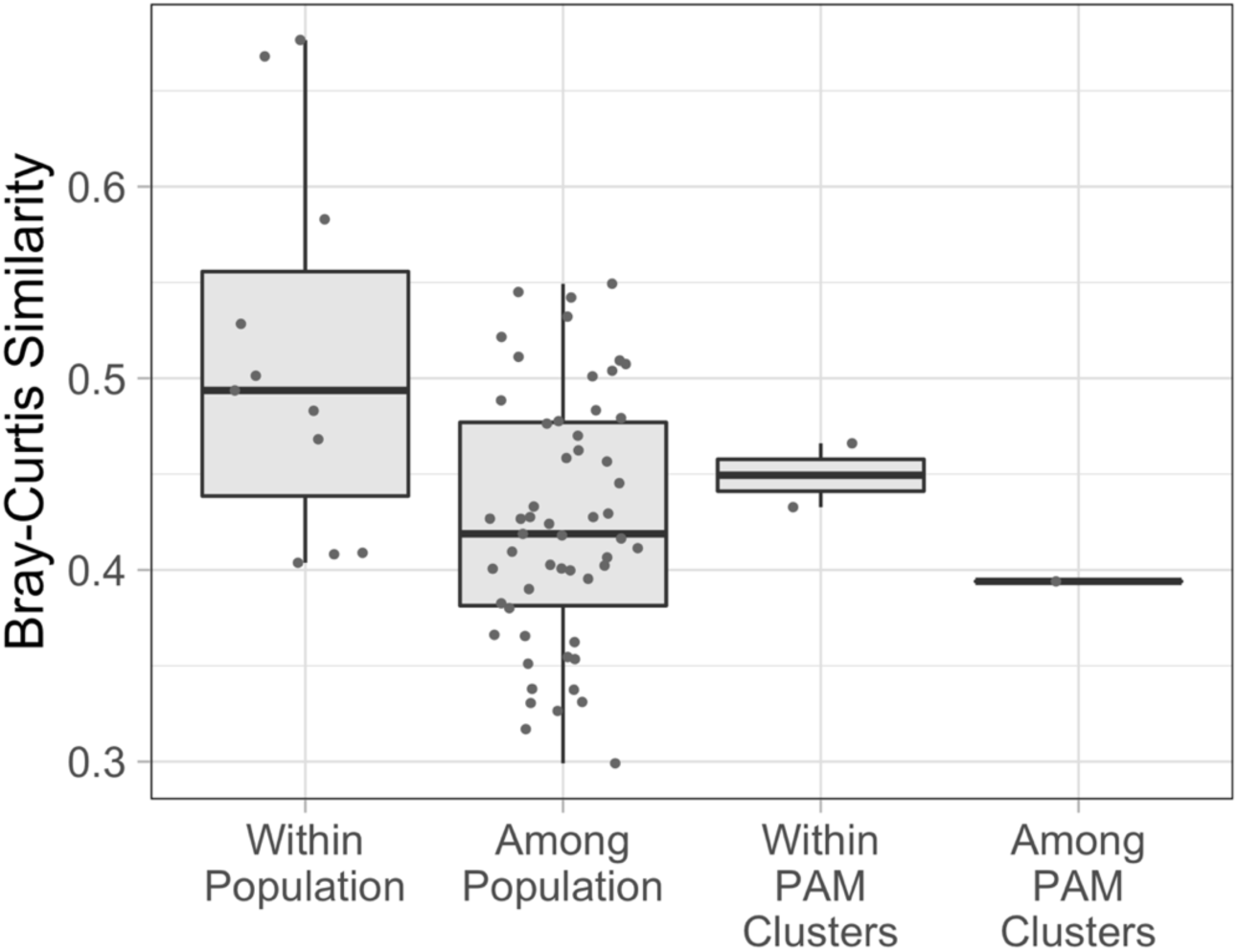
Pairwise Bray-Curtis similarity values from comparing *Arum maculatum* floral odors within and among all sampled populations, and within and among the two PAM clusters identified in this study.

**Figure S3.**
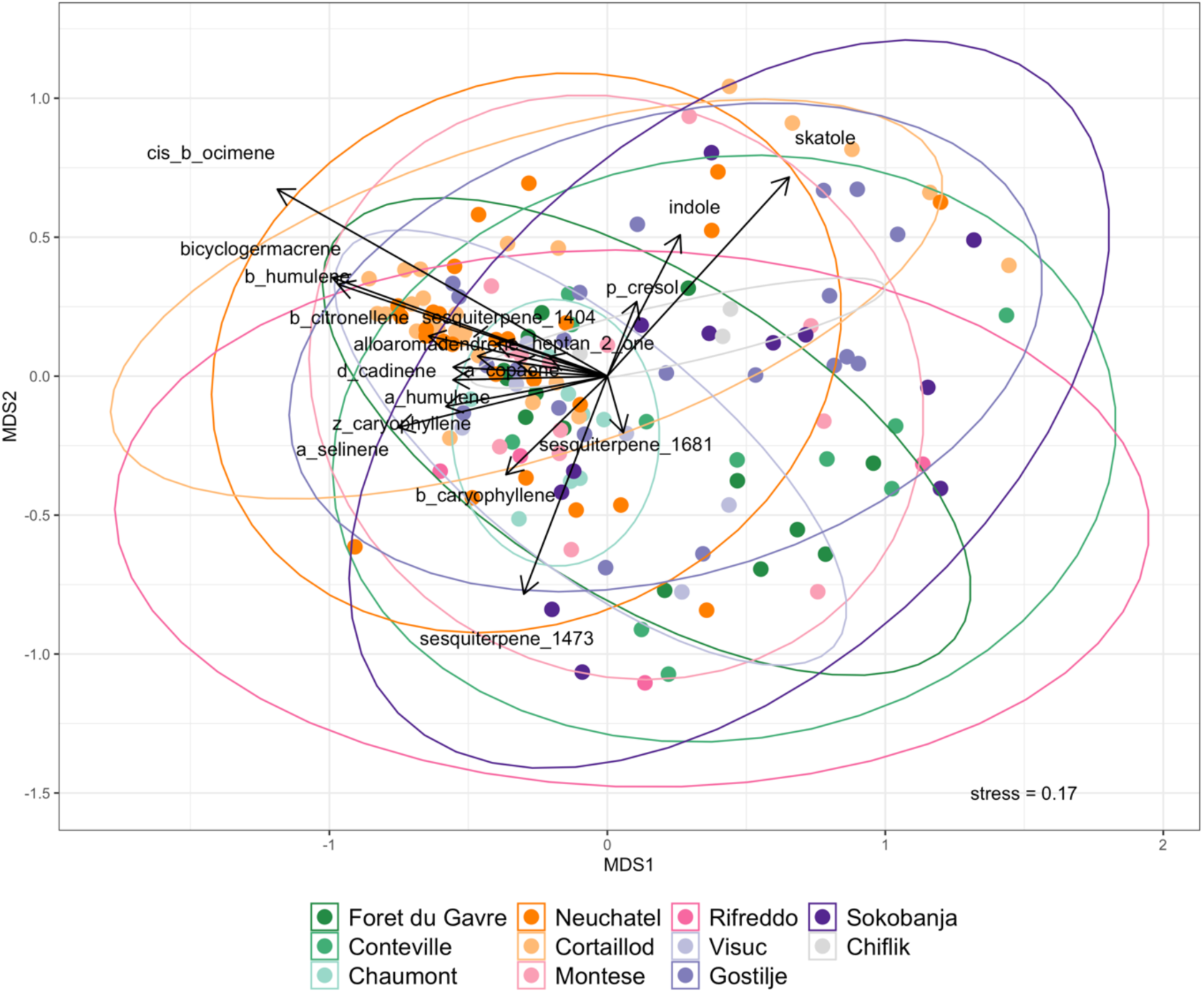
Multivariate representation of *Arum maculatum* volatile organic compound emissions, using nonmetric multidimensional scaling (NMDS) of Bray-Curtis distances between individuals, colored according to their population of origin.

**Figure S4a.**
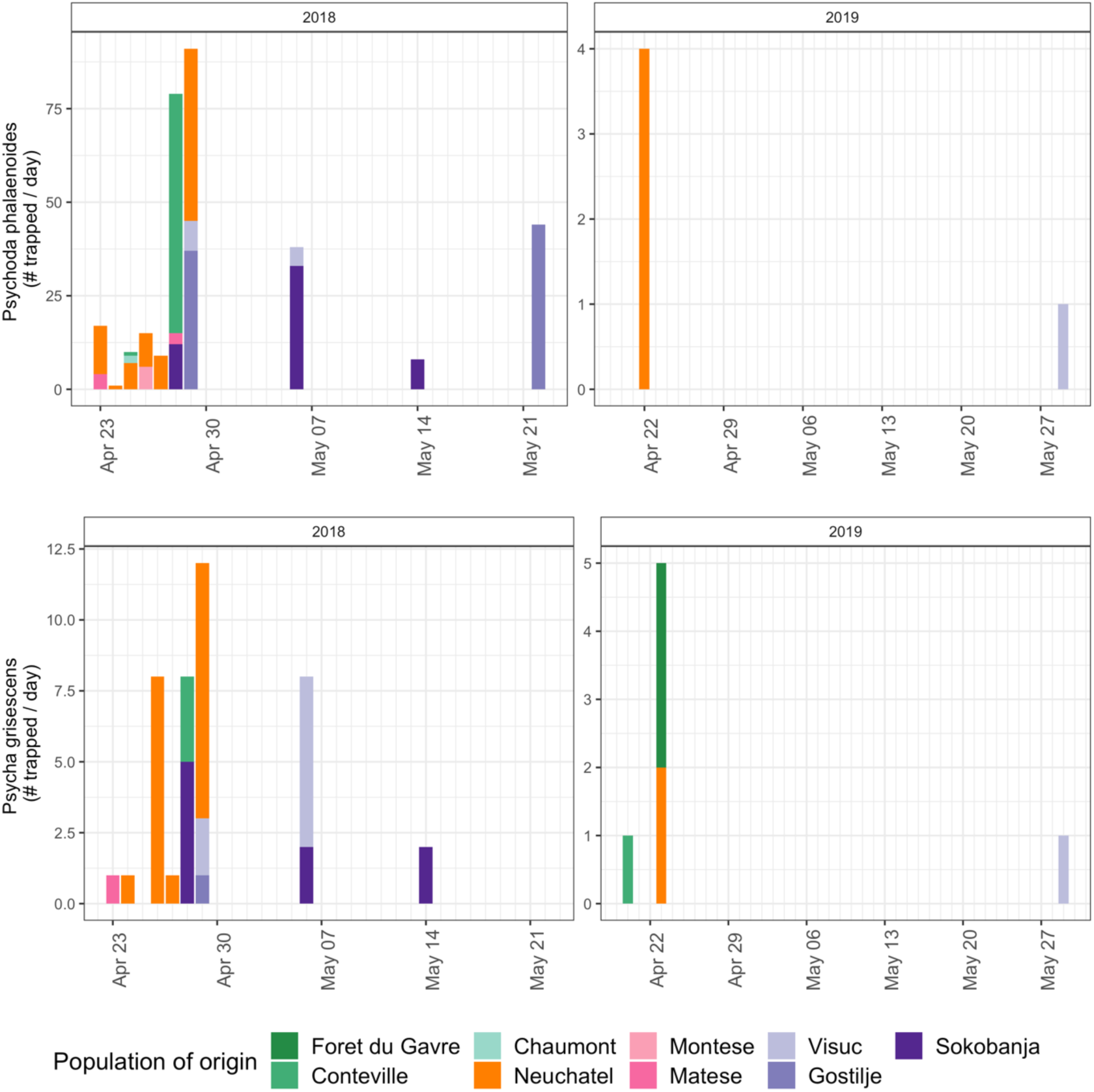
Pollinators trapped by *Arum maculatum* inflorescences in the Neuchâtel common garden (i.e. *Psychoda phalaenoides*-dominated) over two years of sampling. Both *P. phalaenoides* and *P. grisescens* were present and trapped by inflorescences during most of the sampling season. Note: Y-axis scale varies between plots. Bar colors represent the populations of origin for individual inflorescences.

**Figure S4b.**
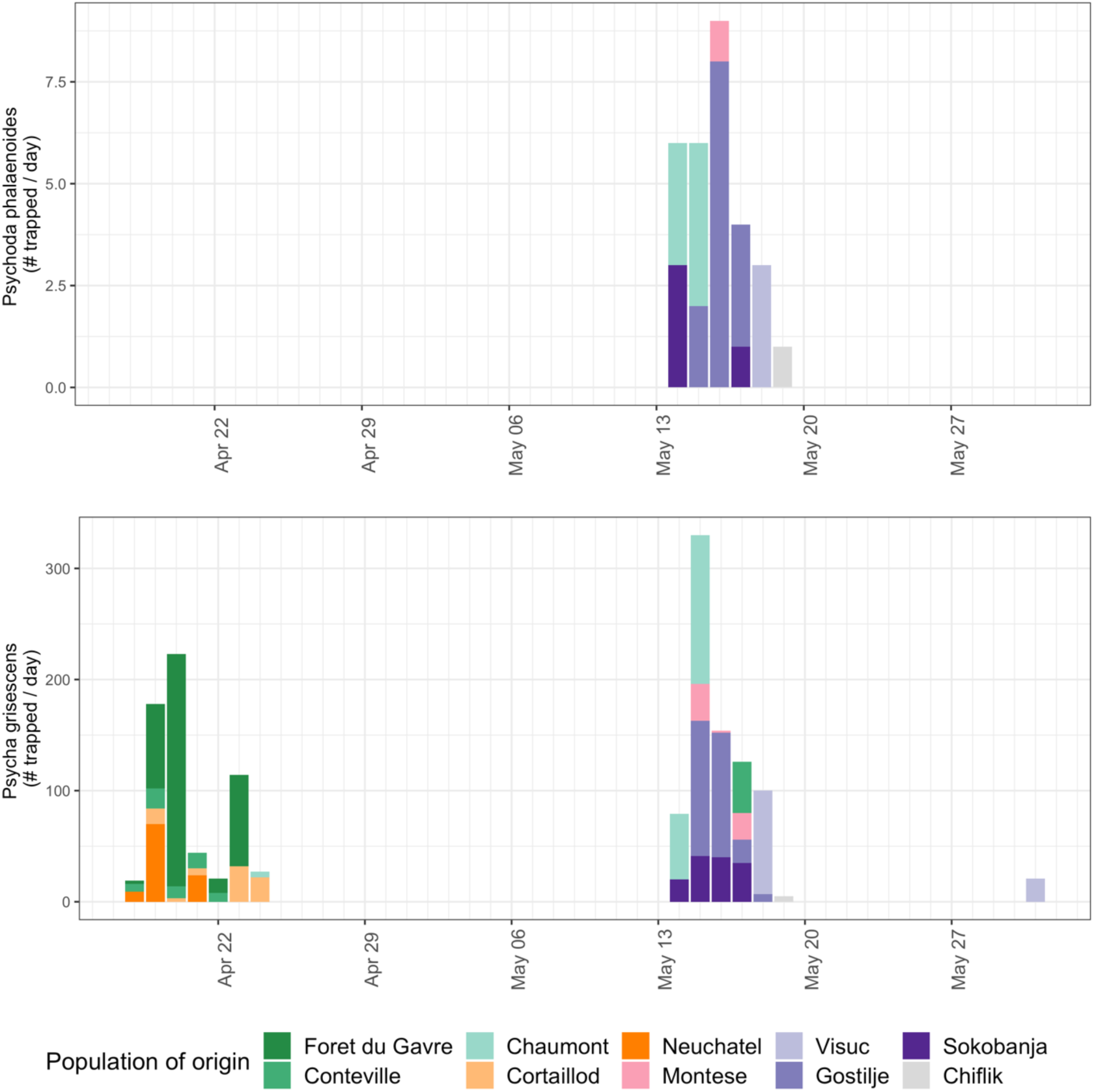
Pollinators trapped by *Arum maculatum* inflorescences in the Forêt du Gâvre common garden (i.e. *Psycha grisescens*-dominated) in 2019. *P. grisescens* were trapped by inflorescences over most of the sampling season, while *P. phalaenoides* were only trapped in the latter half of the season. Note: Y-axis scale varies substantially between plots. Bar colors represent the populations of origin for individual inflorescences. No pollinator sampling was conducted between 25.Apr and 14.May.

**Figure S5.**
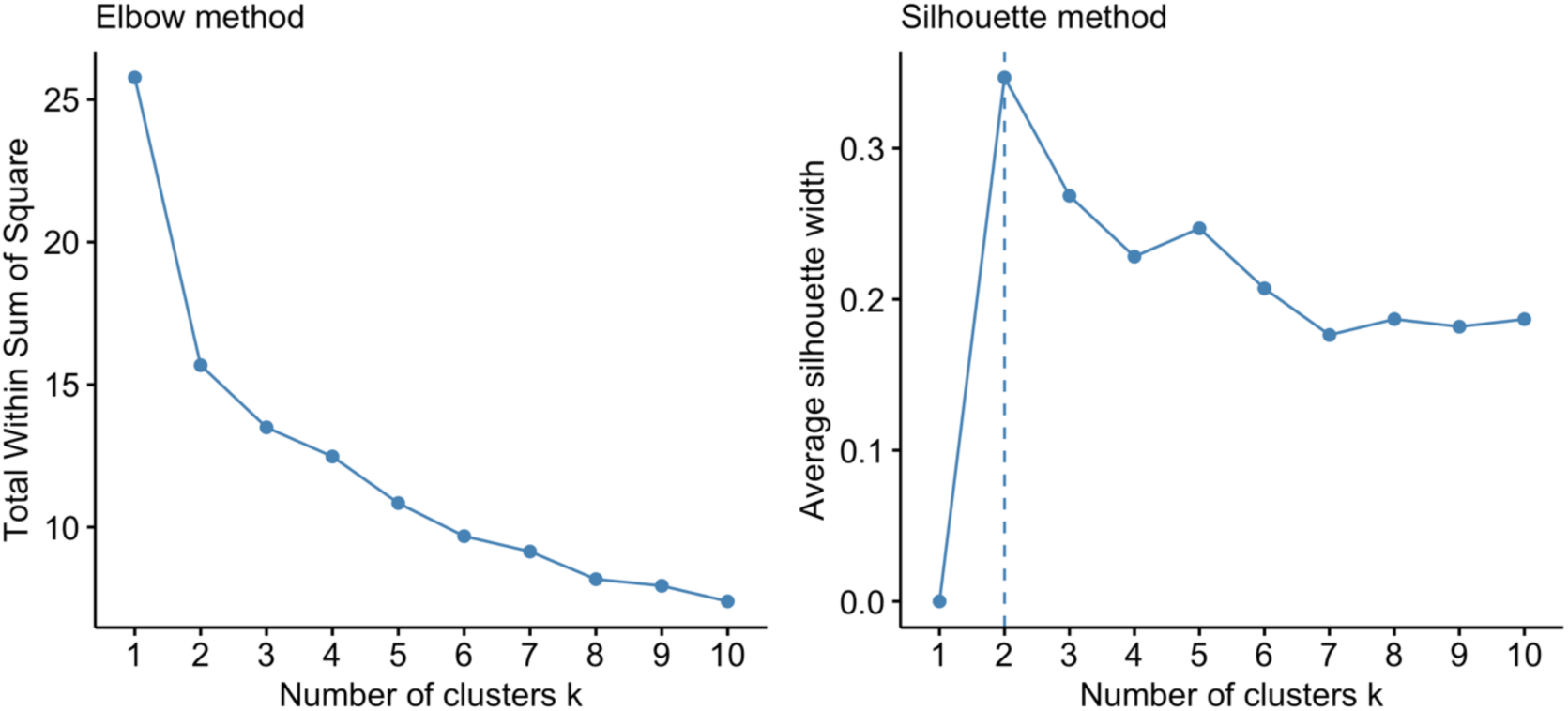
Cluster validation results using ‘Elbow’ and ‘Silhouette’ methods, for clustering of *Arum maculatum* floral odor bouquets into 2-10 groups, using the PAM algorithm. Two clusters appears to be the optimal number for our VOC dataset.

**Figure S6.**
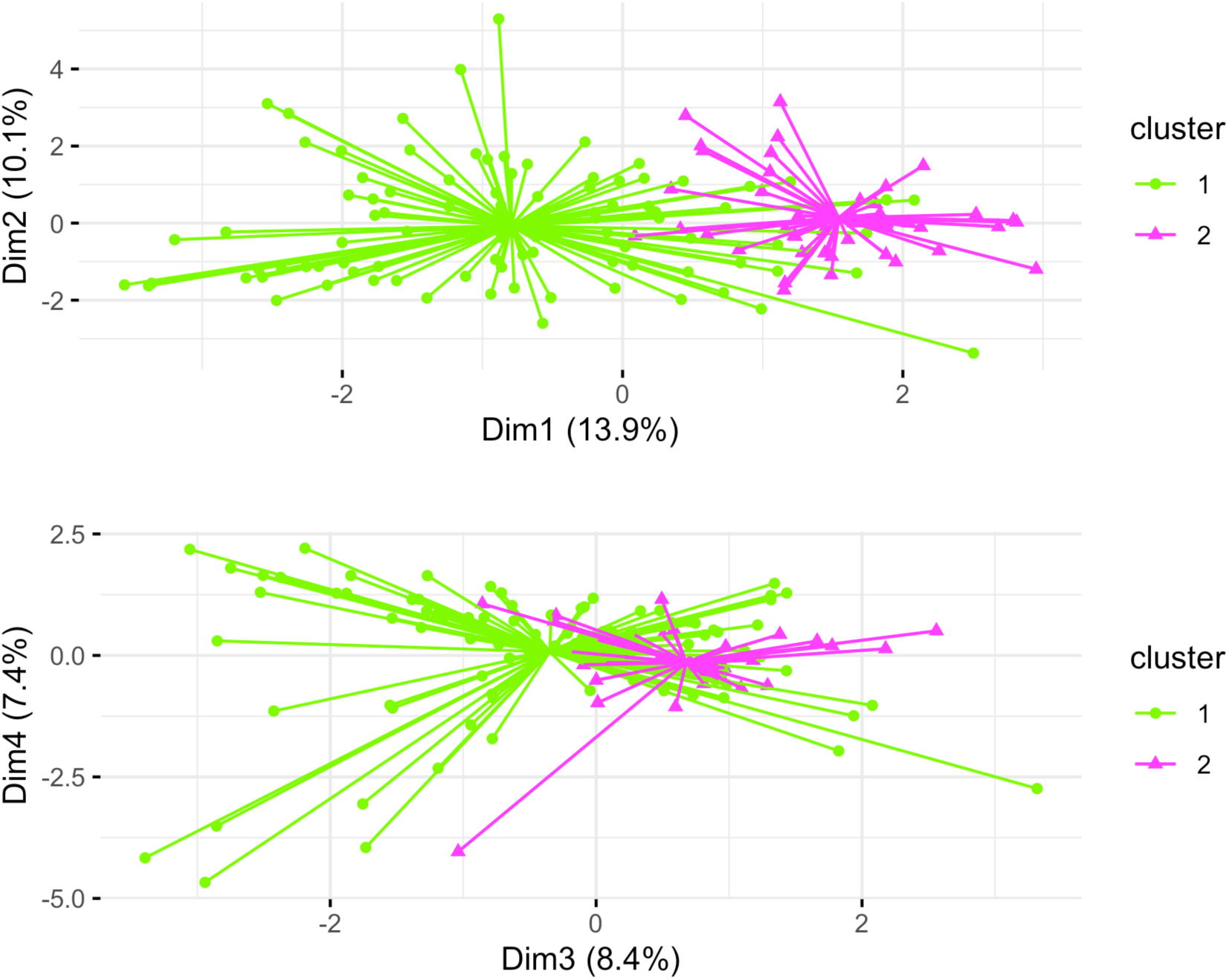
K-medoids (PAM) clustering plots highlighting the placement of *Arum maculatum* individuals into the two clusters we identified, along PCA axes 1 and 2 (upper plot) and 3 and 4 (lower plot).

**Figure S7a.**
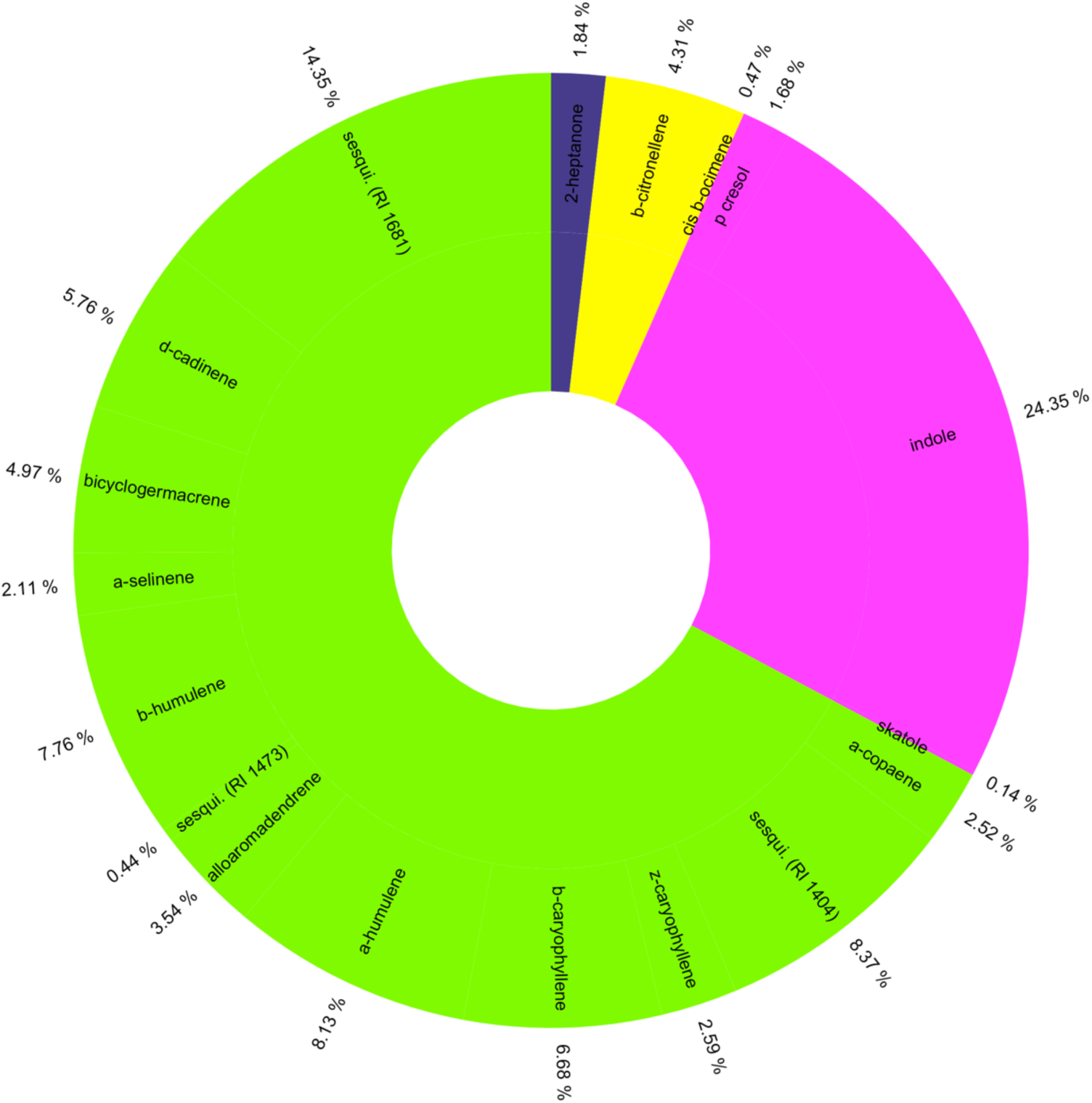
The average composition of VOC blends of *Arum maculatum* inflorescences belonging to PAM Cluster 1. This cluster is mainly characterized by the proportionally large and diverse emissions of mono- and sesquiterpenes.

**Figure S7b.**
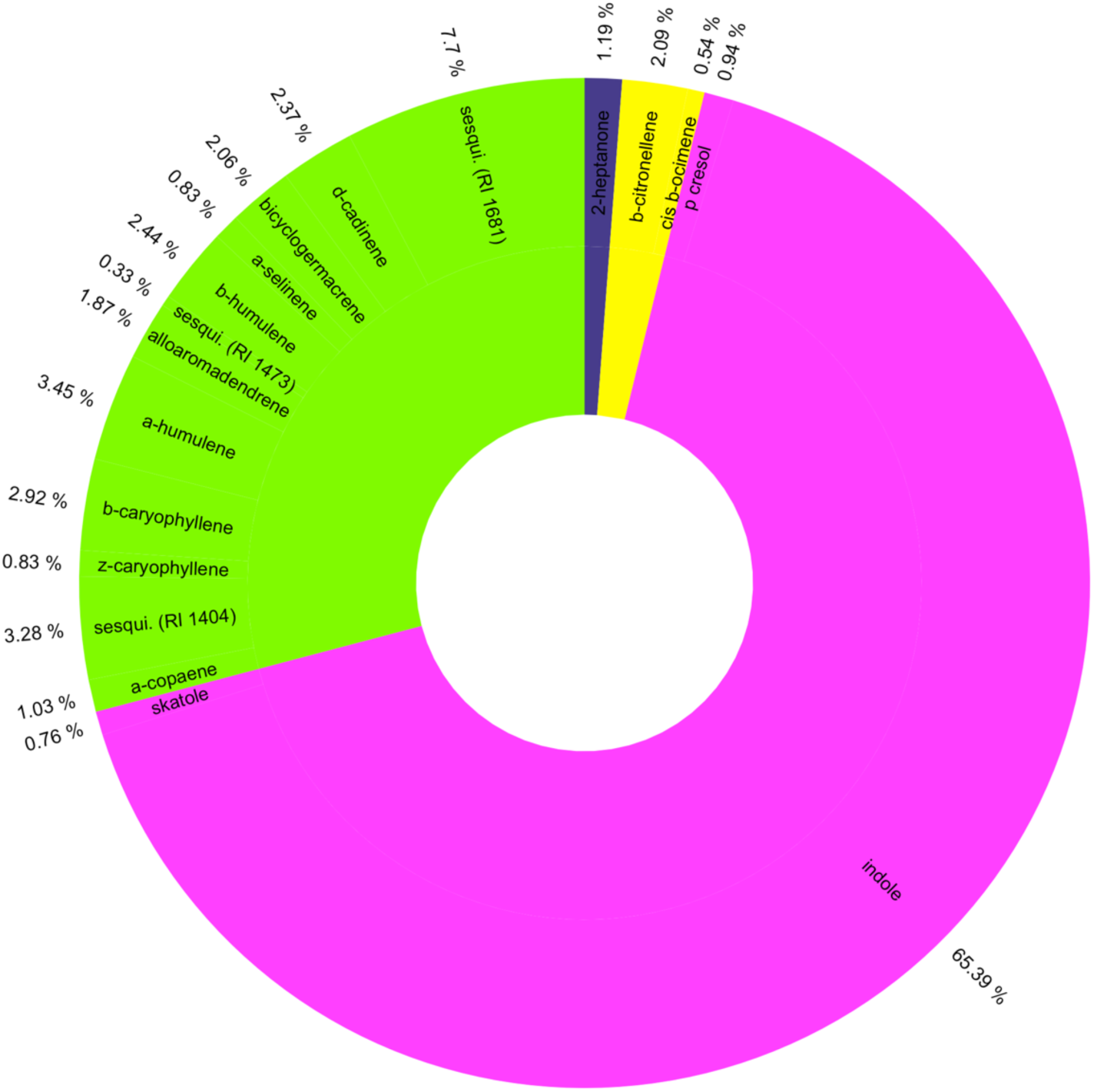
The average composition of VOC blends of *Arum maculatum* inflorescences belonging to PAM Cluster 2. This cluster can be characterized by significantly higher emissions of indole than Cluster 1 (χ2 = 34.92, p < 0.001, df = 1).

**Figure S8.**
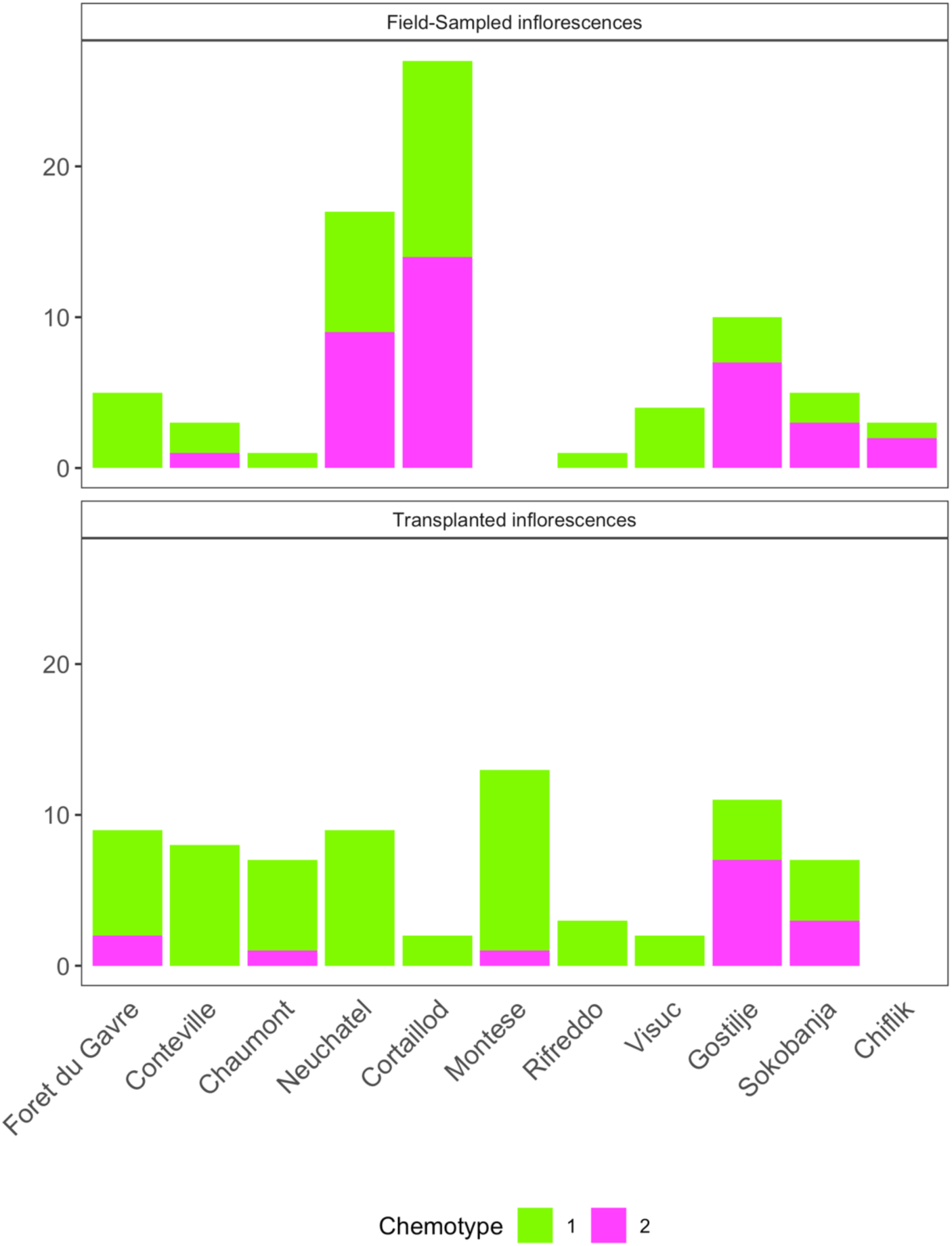
The number of *Arum maculatum* inflorescences assigned to terpenoid-dominated PAM Cluster 1, and indole-dominated PAM Cluster 2. *In situ* VOC samples are shown on the upper graph, while VOC samples from potted and transplanted individuals are shown in the lower graph. No VOCs were able to be sampled *in-situ* in Montese, and following transplants to common gardens for inflorescences from Chiflik.

**Figure S9.**
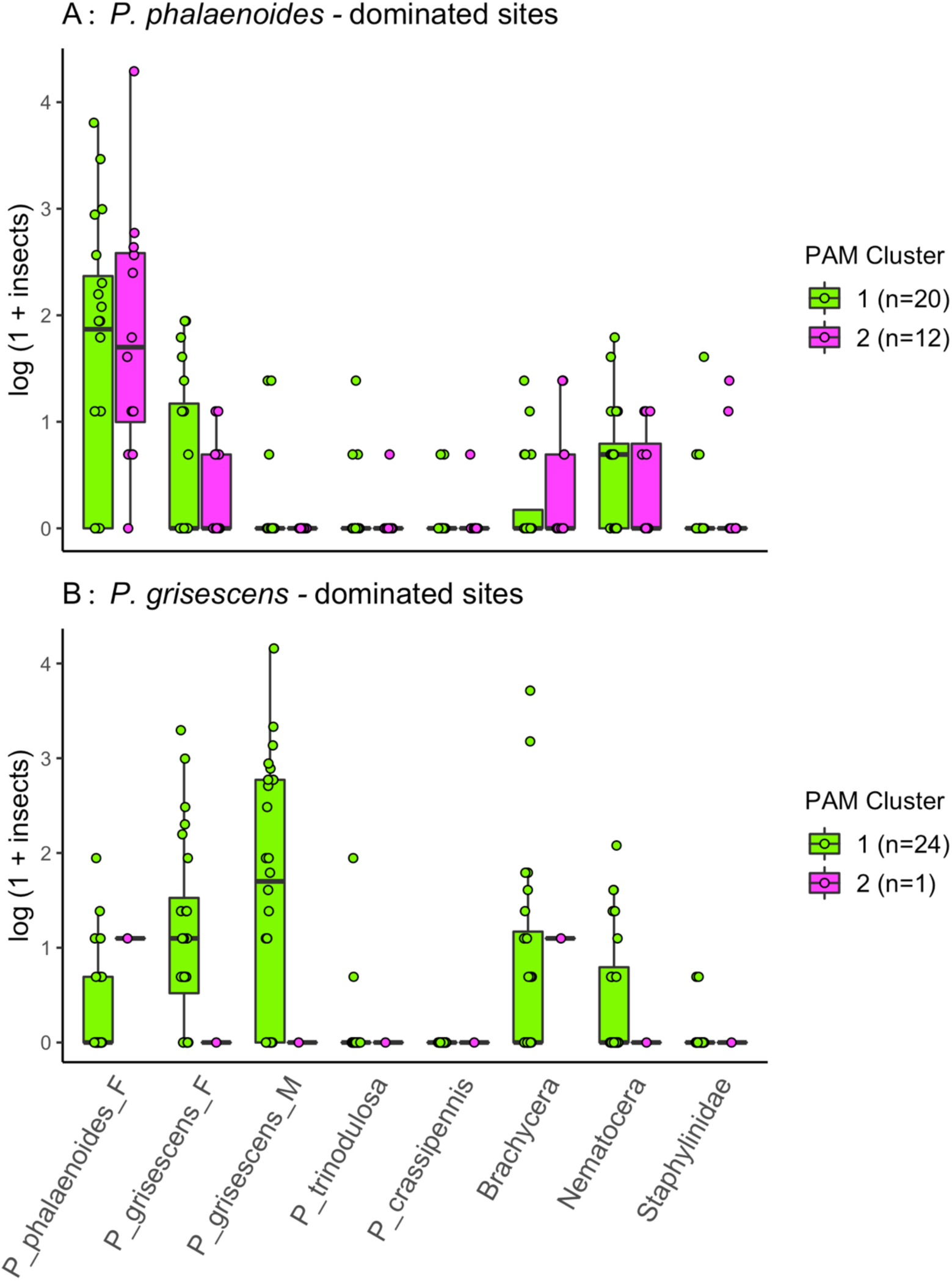
*log1p*-transformed abundances of pollinators trapped by *Arum maculatum* emitting high proportional quantities of terpenoids (PAM Cluster 1) and indole (PAM Cluster 2). To control for geographic variation in background pollinator communities among sampled sites, the visualization is split based on the dominant Psychodidae species where each sample was collected. Where possible, sex (M/F) is specified above; all *P. trinodulosa* and *P. crassipennis* we identified were female.

**Figure S10.**
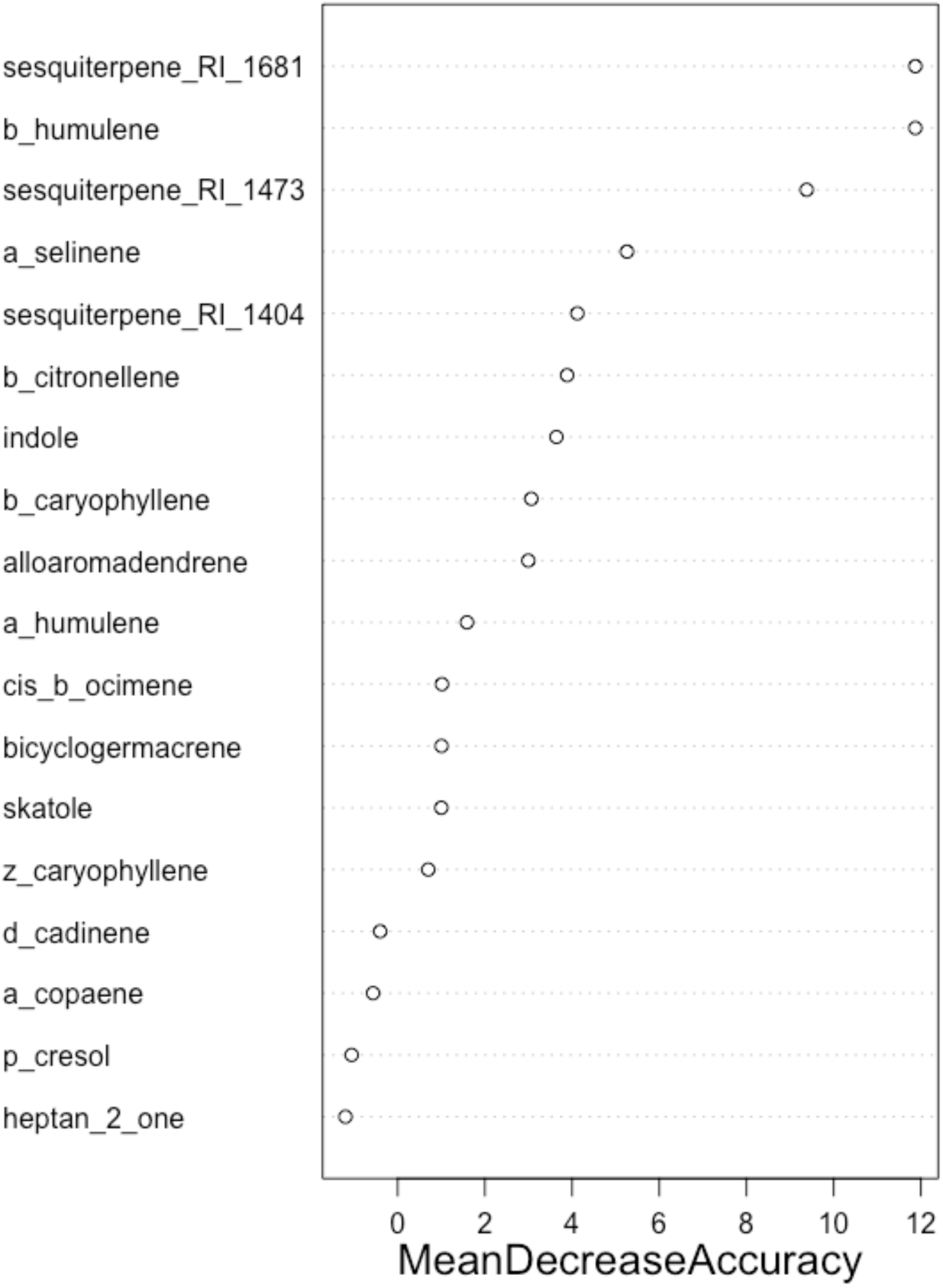
Variable importance plot highlighting the *Arum maculatum* VOCs with the greatest effect on the random forest model accuracy. Compounds such as β-humulene, unnamed sesquiterpenes (RI 1681 and 1473) and, α-selinene were the strongest predictors of whether an inflorescence would trap predominantly *P. phalaenoides* or *P. grisescens*.

**Figure S11.**
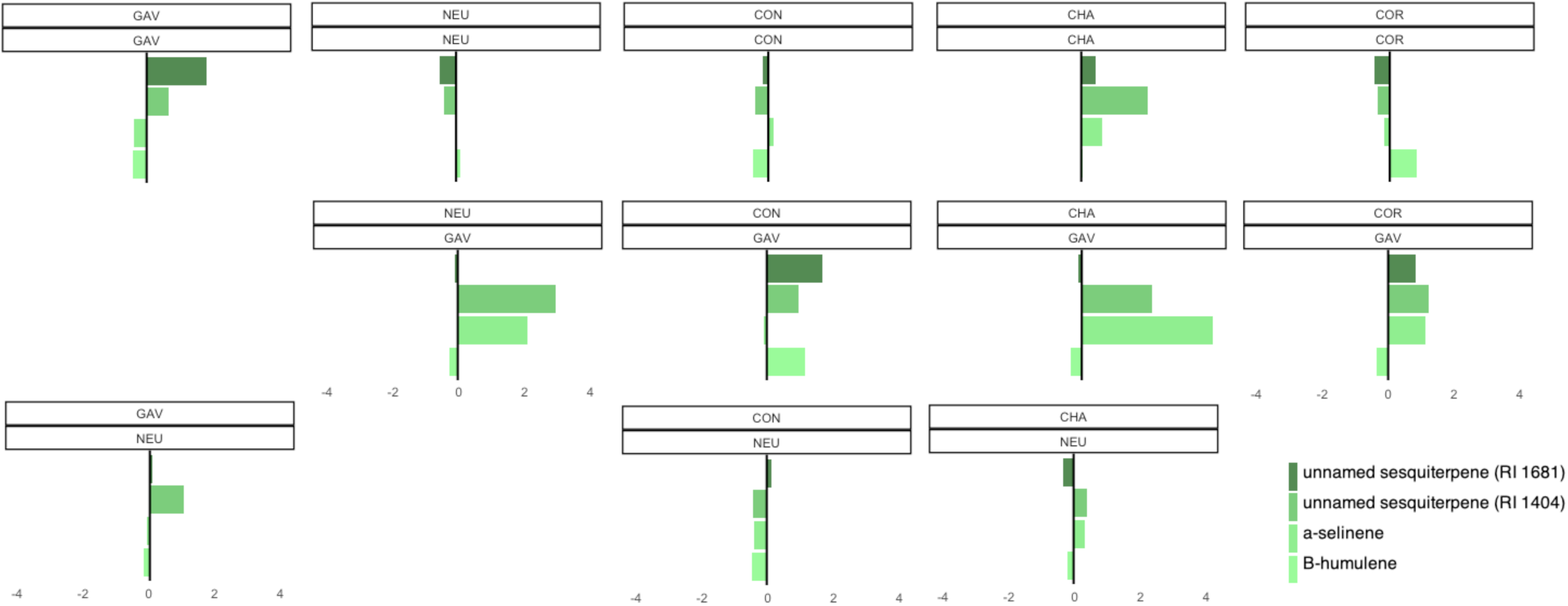
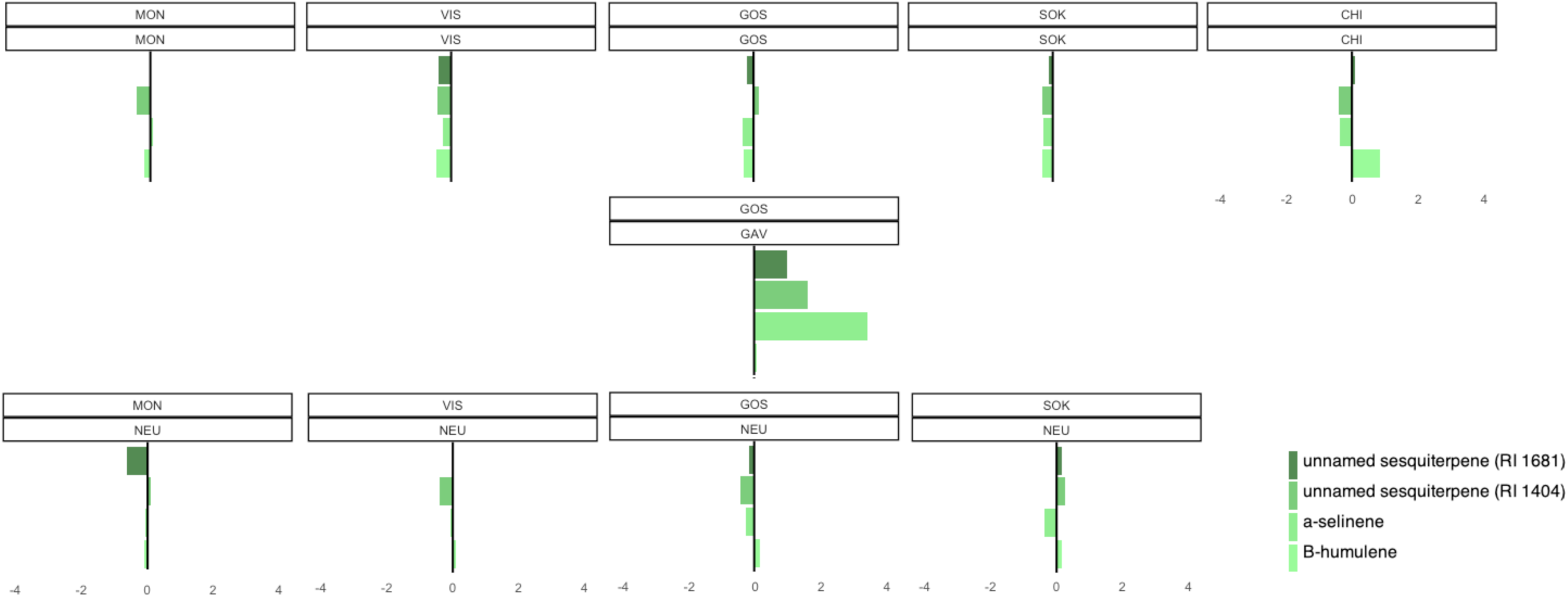
Shifts in the mean population standard scores (calculated as: [(raw individual VOC quantity – mean individual VOC quantity) / VOC std. dev.]) for the four *Arum maculatum* VOCs which were the strongest predictors of whether an inflorescence would trap predominantly *P. phalaenoides* or *P. grisescens*. Two Swiss populations (COR and NEU) and one Serbian population (GOS) shifted to more sesquiterpene-dominated blends following transplants to the Forêt du Gâvre common garden. By contrast, most populations remained relatively consistent in their emissions of these sesquiterpenes following transplants to the Neuchâtel common garden. Note: Upper box represents population origin, lower box represents site where VOCs were collected.

**Table S1.** List of study sites, GPS coordinates, number of Arum *maculatum* VOC samples passing all quality filters, and number of inflorescences with insect data, during both field collections and common garden experiments.

| Population | Country | Lat. | Long. | Alt.<br>ASL<br>(m) | # sampled<br>VOCs<br>(in situ) | # sampled<br>VOCs<br>(NEU,<br>transplant) | # sampled<br>VOCs<br>(GAV,<br>transplant) | # sampled<br>insects<br>(in situ) | # sampled<br>insects<br>(NEU,<br>transplant) | # sampled<br>insects<br>(GAV,<br>transplant) |
| --- | --- | --- | --- | --- | --- | --- | --- | --- | --- | --- |
| Forêt du Gâvre | FR | 47.55066 | -1.86466 | 16 | 10 | 4 | NA | 5 (16) | 3 (3) | NA |
| Conteville | FR | 50.73731 | 1.73872 | 60 | 8 | 2 | 1 | 0 (5) | 1 (4) | 1 (6) |
| Chaumont | FR | 48.11508 | 5.09475 | 296 | 1 | 6 | 2 | 1 (5) | 2 (2) | 1 (5) |
| Neuchatel | CH | 47.00043 | 6.93790 | 556 | 18 | NA | 3 | 8 (30) | NA | 3 (5) |
| Cortailod | CH | 46.93205 | 6.83290 | 430 | 27 | NA | 2 | 9 (14) | NA | 2 (6) |
| Montese | IT | 44.25523 | 10.98371 | 707 | 6 | 6 | 2 | 0 (0) | 1 (1) | 2 (4) |
| Rifreddo | IT | 40.57235 | 15.82473 | 1172 | 4 | 0 | 0 | 0 (0) | 0 (2) | 0 |
| Visuč | HRV | 44.53128 | 15.76134 | 810 | 6 | 2 | 0 | 3 (4) | 2 (4) | 0 (2) |
| Gostilje | SRB | 43.65561 | 19.83549 | 785 | 17 | 3 | 2 | 4 (10) | 3 (4) | 2 (8) |
| Sokobanja | SRB | 43.60373 | 21.88755 | 844 | 10 | 2 | 0 | 1 (11) | 2 (4) | 0 (5) |
| Chiflik | BG | 42.8130 | 24.52836 | 786 | 3 | 0 | 0 | 0 (0) | 0 (0) | 0 (1) |

Note: For the insect data, we report the number of inflorescences with both VOC and insect data (i.e. data used in Figure S9); the adjacent bracketed numbers indicate number of inflorescences with insect data only; this larger sampling was used for all other tests of local adaptation, as well as Figures 2 and 3.

**Table S2.** Mean *in-situ* quantities of *Psychoda phalaenoides* and *Psycha grisescens* caught per *Arum maculatum* inflorescence between 2006 and 2008 (data from Espíndola et al. 2010), and in this study between 2017 and 2019.

| Population | *P. phalaenoides*<br>(Espíndola<br>2010) | *P. grisescens*<br>(Espíndola<br>2010) | *P. phalaenoides*<br>(field surveys<br>2017-2019) | *P. grisescens*<br>(field surveys<br>2017-2019) |
| --- | --- | --- | --- | --- |
| Fôret du Gavre (FR) | 0 (5) | 6.4 (5) | 0 (16) | 25.7 (16) |
| Conteville (FR) | 47.6 (8) | 2.3 (8) | 0.4 (14) | 1.3 (14) |
| Chaumont (FR) | 1.6 (6) | 0.5 (6) | 0 (7) | 0.6 (7) |
| Lausanne/Neuchâtel (CH) | 3.0 (2) | 0 (2) | 2.0 (44) | 0.6 (44) |
| Montese (IT) | 0 (5) | 2 (5) | NA | NA |
| Rifreddo (IT) | 0 (4) | 2.5 (4) | NA | NA |
| Visuc (HRV) | 0.1 (7) | 0.6 (7) | 1.3 (6) | 3.0 (6) |
| Gostilje (SRB) | 0 (5) | 2.4 (5) | 0.4 (18) | 0 (18) |
| Sokobanja (SRB) | 0 (5) | 0.2 (5) | 0 (21) | 0.1 (21) |
| Chiflik (BUL) | 0.4 (7) | 4.3 (7) | NA | NA |

Note: The number of individual inflorescences sampled *in situ* during both sampling periods are indicated in parentheses. NAs indicate that no inflorescences were open yet at the time we visited populations during our study.

**Table S3.** Two-way ANOVA results for pollinators trapped by transplanted *Arum maculatum* growing in two common garden sites, dominated by either *Psychoda phalaenoides* or *Psycha grisescens*.

| ALL POLLINATORS | df | SumSq | MeanSq | F value | Pr(>F) |
| --- | --- | --- | --- | --- | --- |
| common garden loc. | 1 | 2.34 | 2.3369 | 2.138 | 0.153 |
| native poll. | 1 | 0.16 | 0.162 | 0.148 | 0.703 |
| common garden loc : native poll. | 1 | 0.05 | 0.0538 | 0.049 | 0.826 |
| Residuals | 34 | 37.16 | 1.0928 |  |  |

| P. PHALAENOIDES | df | SumSq | MeanSq | F value | Pr(>F) |
| --- | --- | --- | --- | --- | --- |
| common garden loc. | 1 | 18.38 | 18.379 | 16.2 | 0.000301 |
| native poll. | 1 | 0.15 | 0.152 | 0.134 | 0.716174 |
| common garden loc. : native poll. | 1 | 0.01 | 0.008 | 0.007 | 0.931867 |
| Residuals | 34 | 38.57 | 1.134 |  |  |

\*\*\*

| P. GRISESCENS | df | SumSq | MeanSq | F value | Pr(>F) |
| --- | --- | --- | --- | --- | --- |
| common garden loc. | 1 | 36.28 | 36.28 | 55.84 | 1.14E-08 |
| native poll. | 1 | 0.19 | 0.19 | 0.287 | 0.596 |
| common garden loc. : native poll. | 1 | 0.1 | 0.1 | 0.155 | 0.696 |
| Residuals | 34 | 22.09 | 0.65 |  |  |

\*\*\*
**Factors:** **Native pollinator** (whether an inflorescence traps *P. phalaenoides* or *P. grisescens* in their native population) and **Common Garden Location** (Neuchâtel, a site dominated *P. phalaenoides*, or Forêt du Gâvre, a site dominated by *P. grisescens*). F-statistics are shown above (df = 1).
\*\*\* P<0.001, \*\* P<0.01, \* P<0.05

**Table S4.** The top ten combinations of *Arum maculatum* VOCs with the greatest contribution to the predictive strength of the random forest model (see Figure S10). Unnamed sesquiterpene (RI 1681) alone was the strongest predictor of the species composition of pollinators trapped by individual *A. maculatum* inflorescences.

| Compound Blend | Feature Contribution |
| --- | --- |
| unnamed sesquiterpene (RI 1681) | 0.1947778 |
| unnamed sesquiterpene (RI 1473) + unnamed sesquiterpene (RI 1681) | 0.0592275 |
| α-selinene + unnamed sesquiterpene (RI 1681) | 0.0502504 |
| β-humulene | 0.0436112 |
| β-citronellene | 0.0361404 |
| α-selinene | 0.0320748 |
| β-humulene + unnamed sesquiterpene (RI 1681) | 0.0180302 |
| β-citronellene + indole | 0.0138245 |
| unnamed sesquiterpene (RI 1404) + unnamed sesquiterpene (RI 1681) | 0.0119733 |
| unnamed sesquiterpene (RI 1404) | 0.0114971 |

**Table S5.** Mann Whitney U (Wilcoxon rank-sum) test results, comparing the emissions of four VOCs associated with species-specific Psychodidae attraction. To test for shifts in VOCs related to environmental variation, comparisons were made between samples collected in the Forêt du Gâvre common garden and in the Neuchâtel common garden.

| Pop. | α-selinene<br>(*p*) | *Z* | β-humulene<br>(*p*) | *Z* | Sesquiterpene<br>RI 1473 (*p*) | *Z* | Sesquiterpene<br>RI 1681 (*p*) | *Z* |
| --- | --- | --- | --- | --- | --- | --- | --- | --- |
| GAV | 0.0105 | 2.5584 | 0.0213 | -2.3026 | 0.3706 | -0.8954 | 0.1111 | 1.8557 |
| NEU | 0.0102 | 2.5700 | 0.7428 | -0.3281 | **0.0004 \*** | **3.5475** | 0.3128 | 1.0094 |
| CHA | 0.2857 | 1.4652 | 0.8591 | -0.1776 | 0.5942 | 0.5327 | 0.8571 | 0.4637 |
| MON | 1.0000 | 0.0000 | 0.8591 | 0.1776 | 1.0000 | 0.0000 | 0.1798 | 1.3413 |

\* Significant after Bonferroni correction for multiple testing

## Appendix S2 SUPPLEMENTARY FIGURES & TABLES

### Floral odor collection and identification

*A. maculatum* inflorescences open for a duration of roughly 24h, with VOC emissions peaking in the late afternoon / early evening of the first flowering day. We therefore carried out all floral odor sampling between 18:00 at the earliest and 20:30 at the latest. We sampled dynamic headspace volatile organic compounds (VOCs) using polydimethylsiloxane (PDMS) coated Gerstel Twister® (Mülheim an der Ruhr, Germany) stir bar sorptive extraction. Inflorescences were wrapped in inert oven bags (Tangan No34 distributed by Migros, Zurich, Switzerland) cut open at least 8cm above the tip of the spathe to prevent condensation, due to the strong thermogenesis of the appendix. Twisters® were inserted in a glass tube through the oven bag at a height even with, but not contacting, the tip of the spadix. 6L of air was pumped over Twisters® at a standard rate of 200mL per minute for 30 minutes – except for five samples from Conteville, France, where sampling was carried out at the same rate over only 15 minutes. At every sampling site, at least one empty oven bag was placed approximately 5 meters away from any inflorescences, and ambient air was passed over Twisters® identically as with *A. maculatum* inflorescences; these samples were used as controls to filter out ambient air VOCs. All samples were transported in glass vials on ice, and stored at -21°C until analysis.

### Gas Chromatography

We applied 1µL of internal standard (5µg mg/mL naphthalene in dichloromethane) directly to each Twister® immediately before processing. Using a Multipurpose Sampler (Gerstel, Mülheim an der Ruhr, Germany), VOCs were thermally desorbed and separated on a HP-5MS column, 30 m x 0.25mm x 0.25µm at 40°C for 30 sec, increasing temperature by 5°C per min to 160°C, which was held for 0.01 min before increasing 3°C per min to reach 200°C, which was held for 4 min, finally increasing at 10°C per min. until reaching 250°C for 3 min.

### Volatile Data Processing

We aligned peaks by retention time within each population. Major ions were recorded for each integrated peak using Agilent Chemstation software. Putative compound identifications were then derived from NIST 2.3 (library version 17) hits confirmed for the same peak in several spectra; all names used in the final analysis should be considered hypotheses. Compounds present in blank samples with a mean quantity anywhere near those within *A. maculatum* samples were removed prior to further analyses. Quantitative values were obtained by dividing compound peak areas by the internal standard, then multiplying by the internal standard concentration, and finally scaling based on sampling time (for the few samples run for less than 30 minutes).

### Pollinator identification

On the morning after floral odor collection, all trapped pollinators were collected from within each inflorescence and preserved in 70% ethanol. All pollinators were identified to at least the suborder level. Psychodidae were further identified to species level using taxonomic information and illustrations (Ježek 1990). First, the number of antennal segments were counted. 15 segments indicated specimens were likely *P. phalaenoides* or *P. crassipennis*. 16 segments indicated either *P. grisescens* or *P. trinodulosa* -wing venation patterns were then examined for 16-segmented specimens, as *P. trinodulosa* has a characteristic disconnection in one branched vein. To confirm the final species identity (particularly when intact antennae were not available) and sex of all psychodids, the reproductive anatomy of specimens were also examined: Psychodid abdomens were separated, flattened, cleared in a diluted solution of potash, and mounted on a slide in glycerol beside their decapitated head, and wings laid out flat.

### Clustering VOCs using Unsupervised Learning Algorithms

The optimal clustering algorithm (k-means, PAM, or hierarchical clustering) and number of clusters (k = 1 through k = 10) was selected using the clustering validation function implemented in the R package *clValid* (Brock et al. 2008). Following this step, k–medoids (PAM) clustering (Kaufman and Rousseeuw 1990) was used to cluster samples into the optimal number of groups (validated with ‘Silhouette’ and ‘Elbow’ plots); a cluster plot was then created using the R package *factoextra* (Kassambara and Mundt 2019).

We identified significant differences in VOC blend composition among the identified clusters using a Kruskal-Wallis test, and then plotted the mean VOC blend of each cluster. Next, we produced population-level Bray-Curtis similarity matrices, to investigate whether 1) within-cluster similarity was greater than within-population similarity and 2) between-cluster similarity was less than between-population similarity; the variation within each group in the resulting matrices was then visualized using boxplots.

## DATA ACCESSIBILITY STATEMENT

The datasets generated during and/or analysed in this study are not yet publicly available, due to the pending submission of this preprint at a peer-reviewed scientific journal. However, they are available from the corresponding author on reasonable request.

